# Rewiring the Human Brain: On the Fabric of Associative Thinking

**DOI:** 10.64898/2026.04.17.718886

**Authors:** Fabian Czappa, Marvin Kaster, Marcus Kaiser, Xue Chen, Markus Butz-Ostendorf, Felix Wolf

## Abstract

Concept cells are neurons in the medial temporal lobe that represent an abstract concept, such as a familiar person, in a context-independent way. They are activated by heterogeneous and potentially multimodal sensory input related to the concept, such as viewing a photo of the person, reading the person’s name, or hearing the person’s voice. Learning the concept implies connecting the cortical cells that are triggered when such features are perceived with the cells of the concept, resulting in an associative memory engram. Traditional models explain the formation of such an engram with Hebbian learning through synaptic plasticity, the strengthening of existing synapses in response to co-activation. However, the low edge density of the brain suggests that direct connections in the form of preexisting synapses between these cells are relatively unlikely, rendering such constructs inefficient and volatile. Instead, it is plausible that more persistent concept engrams rely on structural plasticity, involving the creation of synapses de novo. Yet, it remains unclear how neurons can find new postsynaptic targets over such a considerable distance. In this paper, we simulate the formation of such structural engrams in the connectomes of healthy subjects, offering a model hypothesis of how such engrams can be formed on a structural level. Based on our model, we further demonstrate how activating a concept can trigger related concepts through overlapping sensory associations, in a process we call a percept-concept loop.

## I. Introduction

Evidence suggests that across primate species, synaptic and neuronal density—the number of neurons and synapses per unit volume—remains roughly constant [1]. This implies that the edge density [2]—defined quite differently as the fraction of possible connections realized in a graph—must decrease as the number of neurons increases because the number of theoretically possible edges grows quadratically with the number of nodes. However, maintaining a constant level of synaptic density is incompatible with a quadratic growth of synapses. The need to avoid excessive space and power requirements—the latter related to signal processing and maintenance—further supports this line of reasoning. The data center of our skull simply cannot host a densely connected brain [3]. In this study, we investigate the functional implications of these structural constraints of network development and reorganization for engram formation and the emergence of conceptual representation in cortical networks.

The more neurons a brain has, the farther its topology is from a fully connected network. For example, the edge density of the human brain is approximately 10^−7^, assuming 10^11^ neurons [4] and 10^4^ synapses per neuron [5]. Brains with large numbers of neurons, therefore, need to make more deliberate choices as to how they wire them; in other words, a lower edge density imposes stricter constraints on the design of their connectome. Like in a computer network, the topology is a tradeoff between price and performance. However, the suitability of a specific network topology depends on the cognitive tasks it is intended to support, which may change significantly over a brain’s lifetime. This provides a strong argument for structural plasticity, which enables the brain to rewire itself to better cope with new challenges. This happens frequently during childhood [6], but also in the adult brain [7], [8]. In general, structural plasticity is assumed to increase the storage capacity of the brain [9].

To stay within the available space and power envelope, new synapses cannot be created at will. Keeping synaptic density below a certain threshold requires a balancing mechanism that offsets the creation of new synapses by discarding those that have become less important. At the same time, this helps keep neuronal activity in a healthy range [10], [11]. Following the same logic, neurons cannot maintain an arbitrary number of active connections to other neurons. Motivated by experimental observations of cortical network reconfiguration after focal retinal lesions in mice, the *model of structural plasticity* (MSP) by Butz and van Ooyen offers such a mechanism [12]. It introduces a homeostatic rule that strives to maintain an equilibrium of neuronal activity. If a neuron’s activity falls below a certain threshold, it starts seeking new connections, encouraging the growth of synapses. If it rises too high, the neuron withdraws from others to evade overstimulation, shedding some of its synapses. Thus, the equilibrium of neuronal activity translates into an equilibrium of synaptic density. In addition, neurons appear to require a minimum level of activity to maintain homeostatic growth, making functional reorganization nontrivial [12].

Building on the initial idea of emulating Hebbian plasticity through homeostatic morphological adaptations of neurons [13], it has recently been demonstrated that structural plasticity can facilitate Hebbian learning [14], [15], which was previously only associated with synaptic plasticity, the weakening or strengthening of existing synapses. This is biologically plausible because synaptic plasticity alone is constrained by the absence of connections. Given the extreme sparsity of the human connectome, missing connections are the rule rather than the exception. Neurons whose spikes can only reach each other by traversing numerous hops can hardly fire together.

Hebbian learning under structural plasticity roughly occurs as follows: Stimulation of selected neurons increases their activity to the point where they begin to prune synapses. Once stimulation ends, they experience input depression, triggering the growth of new synaptic elements, ultimately leading to the formation of new synapses. With a high probability, those will be established between the previously stimulated neurons, providing a network infrastructure that supports their learned association. Driven by the impulse to maintain homeostatic equilibrium, this process derives a significant advantage from its decentralized management. Like in a market economy, rewiring follows demand for and supply of synaptic elements—without the need for superior global knowledge [9].

Yet, it is still unclear how vertebrate brains connect neurons across remote brain areas de novo to bind multimodal perceptual information into new generalized concepts. Concepts such as familiar persons or items are represented by concept cells [16] that start firing when, for example, a familiar person enters the scene. Like a compression algorithm, they integrate only those highly processed percepts, which are indicative of a particular concept but independent of particular declarative episodes [17]. Concept cells are located in the medial temporal lobe (MTL) and require multimodal synaptic input from neurons that encode high-level sensory perception in very different, remote cortical areas [18], [19]. Moreover, the formation of concepts goes relatively fast with only a very few presentations of the corresponding perceptual stimuli being needed to form a stable representation of a concept [20], [21], [22]—in contrast to a more gradual learning in perceptual cortical networks that requires a prolonged training for memorization and successful recall [23]. It is entirely unknown how concept cells establish synaptic contacts with percept-encoding neurons in the cortex to form their individual concept representations. In this study, we employ simulations to demonstrate how structural plasticity can establish new synapses in a highly individualized manner between concept cells in the MTL and percept cells in various cortical regions. Fig. 1 illustrates the underlying concept of associative synapse formation.

**Fig. 1:**
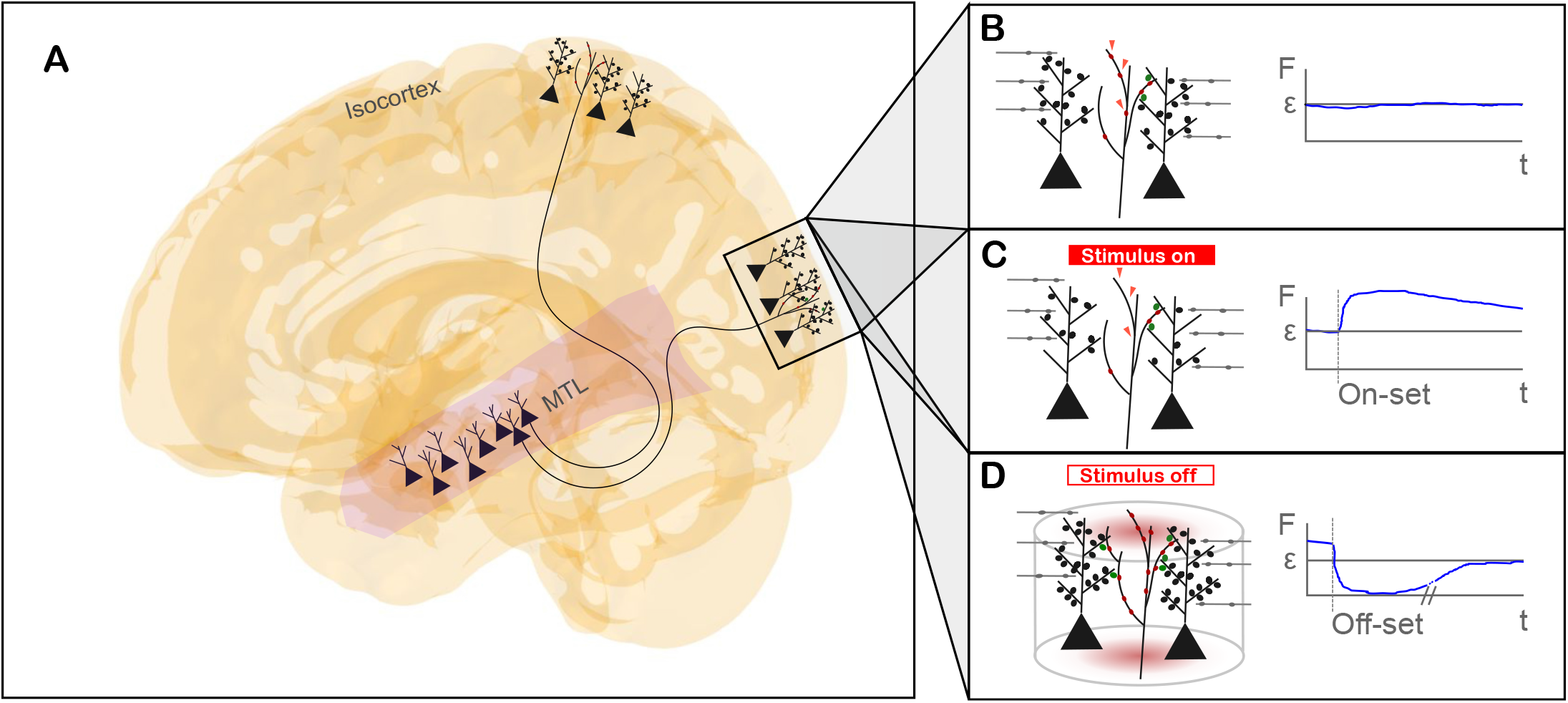
Schematic representation of the formation of engrams in the brain. **A)** Neurons in the medial temporal lobe (MTL) involved in memory encoding send projection fibers to heterogeneous isocortical target neurons processing sensory features. To form lasting memory traces from novel experiences, they need to build synapses between previously unconnected neurons. We illustrate how homeostatic structural plasticity can form synapses in an associative manner without a global observer of spike trains in remote or unconnected neurons. **B)** Before stimulation, activities of neurons in the MTL and in an isocortical target area are in a homeostatic balance. We exemplarily marked some axonal boutons with red arrowheads and followed their fate over time. **C)** In response to synchronous stimulation of MTL neurons and target neurons in the isocortex, firing rates in these neurons increase. They respond with a pruning of axonal boutons and dendritic spines (synaptic elements). Red arrowheads indicate example positions where axonal boutons were eliminated. Loss of synaptic elements can cause synapse breakdown, which decreases firing rates under stimulation, again in a compensatory manner. **D)** Once stimulation is turned off, firing rates drop below initial levels as connectivity was pruned during stimulation. Even previously unconnected neurons in remote locations, sharing the same fate of being deprived of input (via loss of stimulation and pruning of synapses), simultaneously begin to sprout axonal boutons and terminal branches, as well as dendritic spines and branches. These neurons, if in reach (as indicated by the transparent kernel cylinder), have a higher chance to form synapses compared to non-stimulated neurons, which results in associative synapse formation—a direct consequence of homeostatic structural plasticity. Axonal boutons and dendritic spines involved in this form of associative plasticity are labeled red and green, respectively.

This type of learning offers a new perspective on the computations the human brain performs. In the classic model, computation is primarily associated with neural activity driven by spike transmission. Network reconfiguration is seen as a mere act of hyperparameter adjustment, necessary to improve cognitive function. However, as we argue in this paper, structural plasticity can be seen as a computation in its own right, despite occurring on a longer timescale. The connectome, combined with homeostatic plasticity rules, can be interpreted as an automaton—an abstract machine that translates stimulatory input into network changes. Possible network configurations constitute the states of the machine, and the plasticity rules, driven by neuronal activity, define the transitions between them. The initial state is a configuration in equilibrium, subsequently disrupted through a sequence of external stimuli. The final state is the modified configuration reached after the balance has been restored, in a process of consolidation, possibly with minor oscillations around the new equilibrium. The final state then reflects a recallable learned experience, a memory engram.

This paper demonstrates this idea using connectomes derived from living human subjects via brain imaging, made compatible with spiking neuron models and calibrated into a state of homeostatic equilibrium, yielding scaled-down, animatable copies that are ready for learning. Leveraging the structural learning mechanism outlined above, we teach them an abstract concept, resulting in a corresponding memory engram. The engram can be observed by tracing changes in the network. Like one would naturally expect, its precise manifestation is subject-specific. To our knowledge, this is the first simulation of structural plasticity performed on structures extracted from a human brain. In summary, we make the following specific contributions.

- Build large-scale brain simulations of the human cortex based on measured connectomes of healthy subjects in a purely generic manner—without the need for individual parameter fitting.
- Demonstrate that homeostatic structural plasticity can account for Hebbian-style synapse formation between previously unconnected neurons in remote brain regions.
- Simulate the formation of memory engrams in individual human subjects, using concept cells in the medial temporal lobe projecting to co-activated neurons elsewhere in the cortex that represent perceptual features.
- Showcase free memory associations, implemented as a chain of related concepts triggering each other in succession by activating the intersections of their perceptual feature sets.

The remainder of the paper is organized as follows: Section II presents our results, which Section III discusses in a broader context. Section IV details our methods, while the appendix provides supplementary material.

## II. Results

In theoretical neuroscience, learning has traditionally been equated with synaptic plasticity, most notably with Hebbian learning [24]. Classical mechanisms such as long-term potentiation (LTP) and long-term depression (LTD) [25] exemplify this approach and have extensive applications in theory and modeling [26], [27]. In spike-timing-dependent plasticity (STDP) [28], for instance, synapses strengthen when the postsynaptic neuron fires shortly after the presynaptic neuron. Conversely, the opposite timing leads to synaptic weakening. While this framework captures essential aspects of activity-dependent learning, it oversimplifies the complex dynamics present in real neural networks.

Recent studies [14], [15] suggest that homeostatic structural plasticity [12], [29] can produce similar associative effects without explicitly modeling synaptic plasticity. In these models, learning emerges from self-regulating mechanisms that adjust neuronal firing rates toward a set-point. Through the interaction of such homeostatic regulation across a network, correlations in activity can be reinforced, effectively mimicking Hebbian associations. This perspective challenges the classical view that synapse-specific plasticity is strictly necessary for associative learning. Instead, it emphasizes the potential role of network-level, homeostatic processes in shaping neural representations. Building on these insights, we simulated associative learning in the connectomes of healthy subjects. The overall workflow is summarized in Fig. 2. We obtained the connectome datasets from the control group of an epilepsy study [30] via diffusion tensor imaging (DTI), cortical triangulation, and fiber-tract reconstruction. This resulted in neural networks with approximately 47 000 nodes per subject. Next, we transformed these networks into animatable *avatar* connectomes. We interpreted their nodes as neurons and their edges as synapses in a process we call *neuronization*. Then, we brought them into a state of homeostatic equilibrium to make them susceptible to learning in a process we call *homeostatization*. We subjected the homeostatized connectomes to a stimulation protocol that taught them abstract concepts by linking neurons in the MTL to cortical regions assumed to host representations of sensory input. Synapses were rewired according to an extended version of the MSP rule set that largely preserves biological axon-length distributions by always selecting target neurons within the immediate vicinity of the soma or the telodendron of the axon. In a recall experiment, we validated learning success and explored association chains by activating related concepts via overlapping perceptual contexts. Below, we describe our results in detail and demonstrate our ability to reproduce Hebbian-style learning phenomena in humans solely on the basis of structure. Beyond what is reported here, further details on our imaging pipeline, the brain model, and MSP can be found in Section IV.

**Fig. 2:**
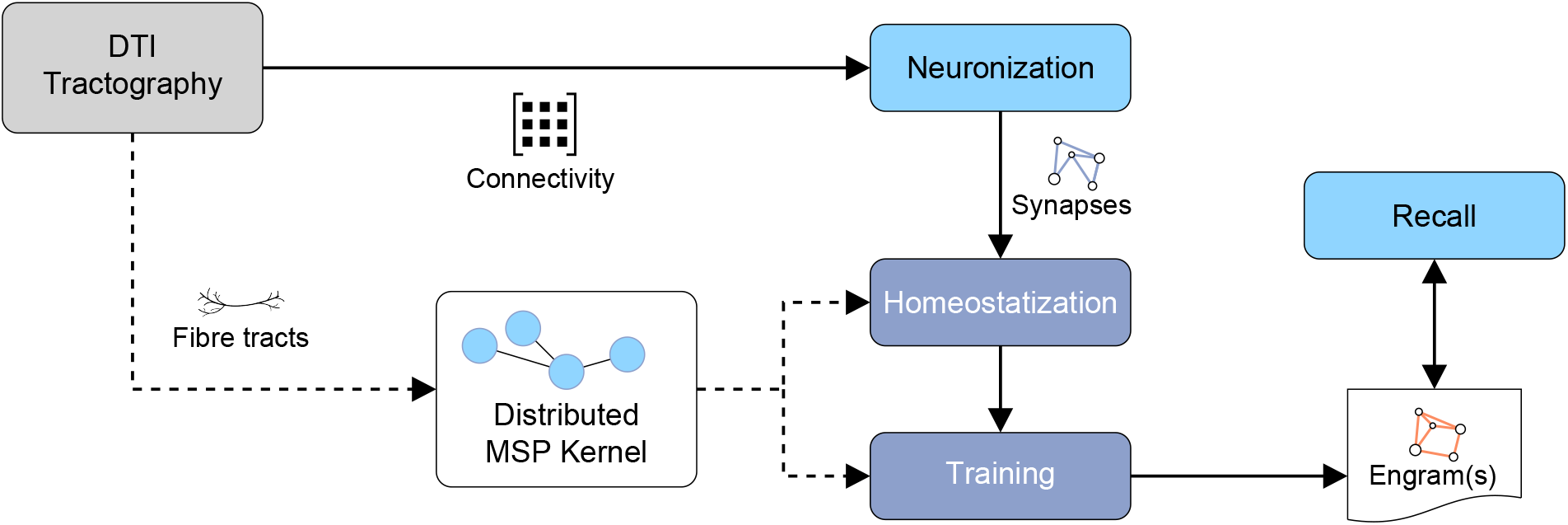
Starting from a connectivity matrix obtained using DTI tractography, our workflow consists of four phases (blue and gray boxes): (1) We interpret the nodes of the connectome as neurons and the edges as synapses. This process is referred to as neuronization. (2) We bring the connectome into homeostatic equilibrium to prepare it for learning, a process called homeostatization. (3) We train the connectome to teach it abstract concepts, resulting in one or more engrams. (4) Finally, we perform a recall experiment to activate the learned concepts via their engrams. Steps 2 and 3 modify the connectivity, with the fiber tracts serving as guardrails enforced by an extended version of the MSP probability kernel. The tracts themselves remain unchanged throughout the workflow.

### A. Neuronization

Obtaining structural connectomes of the human brain is challenging. We define the *structural connectome* as a directed graph of synaptic connections between neurons, from the presynaptic to the postsynaptic neuron, disregarding axonal and dendritic morphology and synaptic strength for now. Brain imaging techniques, such as DTI for live subjects and polarized light imaging (PLI) [31] for deceased subjects, fall short of extracting the true graph. First, these techniques capture fiber tracts rather than synapses themselves. Metaphorically speaking, we see the ductwork system rather than the cables running through it. Consequently, we can only make educated guesses about where a synapse originates or terminates. Second, these techniques operate at resolutions far lower than the size of neurons and synapses requires. Although imaging technology has improved significantly over the past few decades [32], it remains uncertain whether it will ever be possible to obtain an accurate connectome of the entire human brain. Microscopy, on the other hand, can only analyze small tissue samples. For the time being, therefore, such connectomes will likely remain approximations. Although the resolution of the connectomes used in this study is high by today’s standards, it is six orders of magnitude lower than that of the human brain. This is akin to a map of highways and major country roads that omits streets and narrow alleys in residential areas. Additionally, their node-degree and edge-length distributions exhibit highly irregular patterns that differ significantly from those observed in tracing or microscopy.

So, how can we simulate the human brain when precise biological network data is lacking? How can we give virtual life to the structures obtained from brain imaging to better understand their functional properties? To enable simulations based on approximate connectomes, we neuronize them: We interpret the nodes of the measured connectome as single neurons, even though they are actually groups of neurons, and the edges as synapses between those neurons. This is an alternative to the neuron-mass model employed in The Virtual Brain [33], which focuses on the average behavior of the neuron population a node represents, such as the mean membrane potential and average firing rate. The primary benefit of our approach is that it allows us to incorporate spiking neuron models and MSP, a neuron-level model for structural plasticity. Because our imaging pipeline does not produce directed graphs, we further translate each unidirectional edge into two synapses, one in each direction.

### B. Homeostatization

Moreover, we calibrate the connectome to improve its functionality and biological plausibility. This is achieved by moving it into a state of homeostatic equilibrium. Imaging artifacts, primarily an unnaturally imbalanced node degree distribution, render raw connectomes unsuitable for learning with MSP because the imbalance leads to inhomogeneous activity that distracts neurons from forming connections between co-stimulated assemblies, much like classroom noise makes teaching ineffective. Additionally, the edge-length distribution exhibits unexpected anomalies. Homeostatization smooths the node degrees and aligns the axon length distribution with known estimates [2]. A major advantage of our method is its self-regulating nature. It achieves the desired transformation without requiring adjustments to its underlying model parameters.

The changes to the topology are neither insignificant nor profound. We compared the connectomes before and after homeostatization using axon length and node degree distributions (Fig. A5) and complex network measures (Fig. A6). The distribution of axon lengths stayed fat-tailed, maintaining long-range connections. Consequently, characteristic path lengths, global efficiency, and total synapse numbers remained almost unchanged. Although the clustering coefficient decreased following local connectivity refinements, the small-world property of the homeostatized connectomes stayed intact. Overall, the physiological network’s characteristics, such as clustering, were well preserved. Homeostatization maintained efficient communication through long-range connections, with the necessary rearrangements occurring predominantly in the nodes’ in- and out-degrees. Essentially, homeostatization is like remastering a soundtrack, eliminating noise and adjusting the balance between quiet and loud sections. It improves the experience, but neither alters the composition nor the arrangement.

The combined result of neuronization and homeostatization is a scaled-down version of a living person’s connectome, a miniaturized avatar connectome that can be animated to study learning and memory. Note that the need for miniaturization (i.e., having far fewer neurons than a real brain) is not a limitation of our approach but rather of the imaging pipeline, which cannot yet achieve neuron-level resolution. Our approach, therefore, bridges today’s scale gap between imaging and neuron-level simulation.

### C. Training

Since our model neurons correspond to the nodes obtained from tractography and inherit their 3D coordinates, our brain model encompasses the entire cortical surface. There, each neuron is mapped to one brain area defined by the Desikan–Killiany atlas [34]. Here, we focus on the MTL because it contains single cells that bind together different sensory input modalities. Examples for such cells include place cells [35], memory cells [36], schema cells [37], and concept cells [38]. Although the resolution of the avatar connectome is too low to distinguish different cell types, we may assume that each triangle of the triangulated MTL surface likely contains a concept cell next to a variety of other cell types. We therefore define that, after neuronization, a subset of neurons in the MTL shall represent *concept cells* (Fig. 3a). We model concept cells for two reasons: First, forming novel concepts requires binding very different and unpredictable sensory features. Second, this binding occurs quickly based on only one or a few stimulus representations. These properties are difficult to achieve with traditional forms of Hebbian learning, such as STDP. Throughout the cortex, principal neurons— here called *percept cells* (Fig. 3b)—encode part of the multimodal sensory perception of a familiar person, for example, to which the group of concept cells responds. For simplicity, our probability kernel permits synapse formation directly between concept cells and their associated percept cells. In reality, a hierarchy of memory-encoding cells exists in the MTL, and connections to distant sensory cortices may pass through relay neurons; our simplification does not compromise the model’s validity, however, since the plasticity rule applies equally to all cell types involved. Thus, our model predicts the localization of engrams forming in an individual’s brain.

**Fig. 3:**
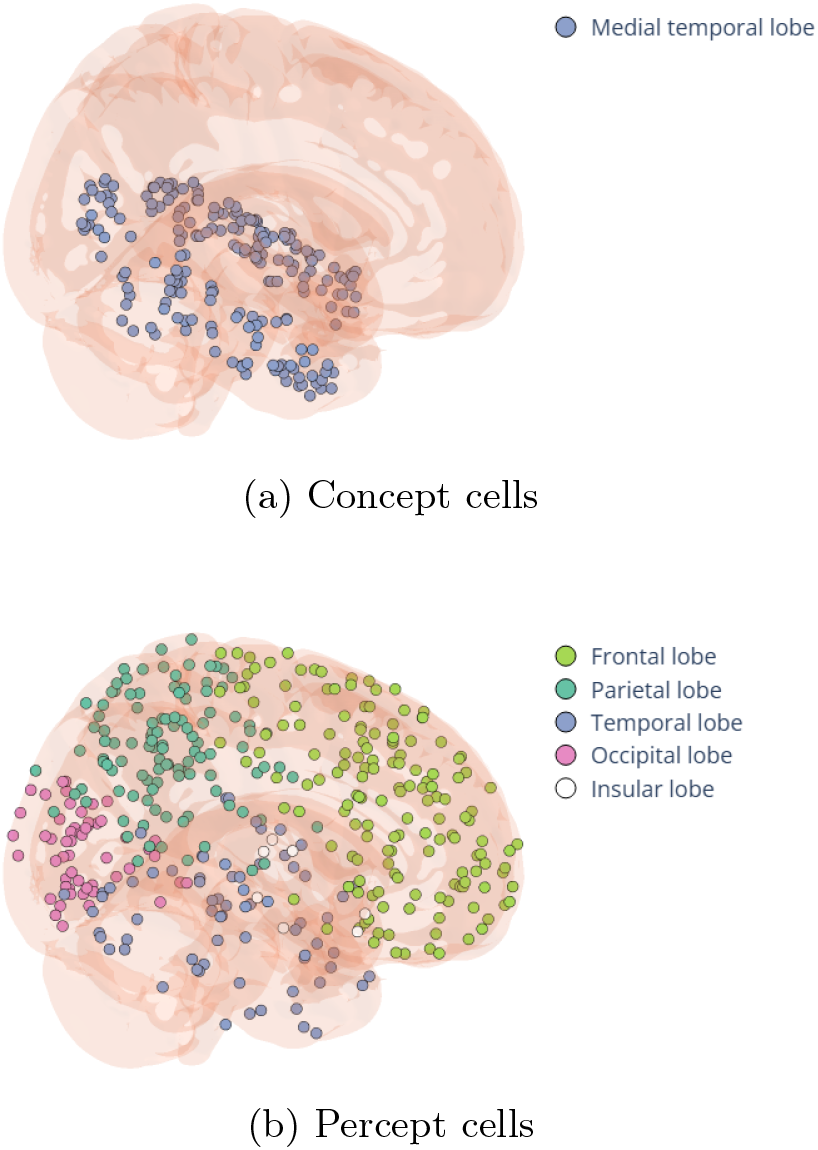
Designated neurons for one engram, divided into 200 concept and 400 percept cells.

An *engram* is a memory trace that leaves lasting structural and functional changes in brain networks in response to a novel experience [39], [40]. We designated 200 MTL neurons, a small subset of the MTL neuron population, as an assembly of concept cells (or neurons). We use the acronym CC to denote such an assembly, which can become the physical representation of a learned concept. Additionally, we selected twice as many neurons distributed across the cortex outside the MTL as percept cells to represent highly processed sensory features, such as faces, voices, objects, and places. We use the acronym PC to denote an assembly of percept cells, which can become the physical representation of a percept—the collection of perceptual features linked to a concept. Fig. 3 displays the location of the two assemblies.

Next, we investigated whether stimulating a CC and a PC simultaneously would promote synapse formation within each group and between them, potentially across different brain areas. We define a *structural engram* as the set of newly formed synapses between a CC and a PC that were absent before training. To induce memory engrams, we applied the stimulation protocol in Table A3a. This protocol, adapted from previous work [15], was motivated by evidence that brain-wide spiking activity aligns with MTL slow waves to support human memory consolidation [41]. The training protocol was applied after homeostatization. It consists of two phases: (i) stimulating the CC and the PC individually and (ii) stimulating them together. The first step strengthens the internal cohesion of each assembly, establishing them as groups. The second step increases the cohesion between these groups. Fig. 4 illustrates the stimulation timeline and its effects. It exposes how existing synapses retracted and new ones formed, ultimately leading to engram formation.

**Fig. 4:**
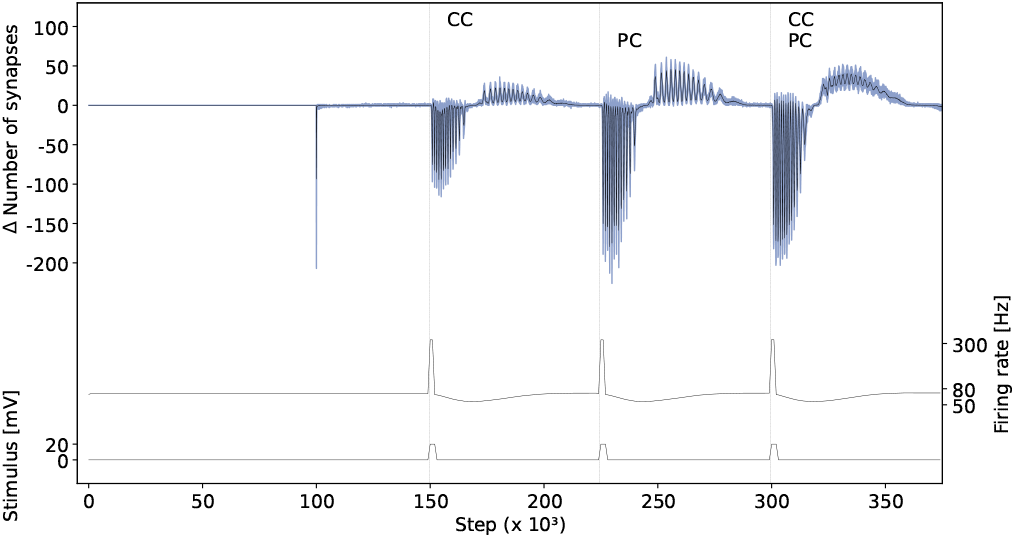
Influence of training on neural activity and the retraction and growth of synapses. According to our stimulation protocol in Table A3a, training a single engram involves three distinct stimulation events (bottom of this figure); first, the concept-cell assembly (CC) and the percept-cell assembly (PC) are stimulated in isolation. Then, they are stimulated together. The assembly identifiers are displayed at the very top. Each event causes the neural activity (i.e., firing rate) in the stimulated assemblies to peak, followed by a wider but shallower slump (middle). In response to activity exceeding the set-point, neurons shed existing synapses to compensate for their increased activity (top). During the subsequent period of low activity, neurons that had previously lost synapses formed new ones until the original activity level was restored. Because all of the co-stimulated neurons were seeking new partners at the same time, they tended to connect with each other. This resulted in the formation of the engram. The blue region represents the spread of the difference in the number of synapses relative to the previous time step across all 36 connectomes, while the black line represents the average. This number reflects changes throughout the entire connectome, including neurons outside of the stimulated assemblies. Since the stimulation protocol was identical across connectomes, and the networks were nearly identical in size with the same number of stimulated neurons, the traces from the individual connectomes overlap almost completely.

Fig. 5 shows the relevant connectivity features of the structural connectome before and after homeostatization and training, averaged across all 36 connectomes. Before training, CC neurons were primarily connected to neurons that were neither concept nor percept cells. After training, intra-CC connectivity increased tenfold on average, while inter-area connectivity from CC to PC and vice versa increased by factors of around 40. The success of the training depends critically on achieving homeostasis beforehand (Fig. 5d). Fig. 6 displays the synapses created for one specific connectome.

**Fig. 5:**
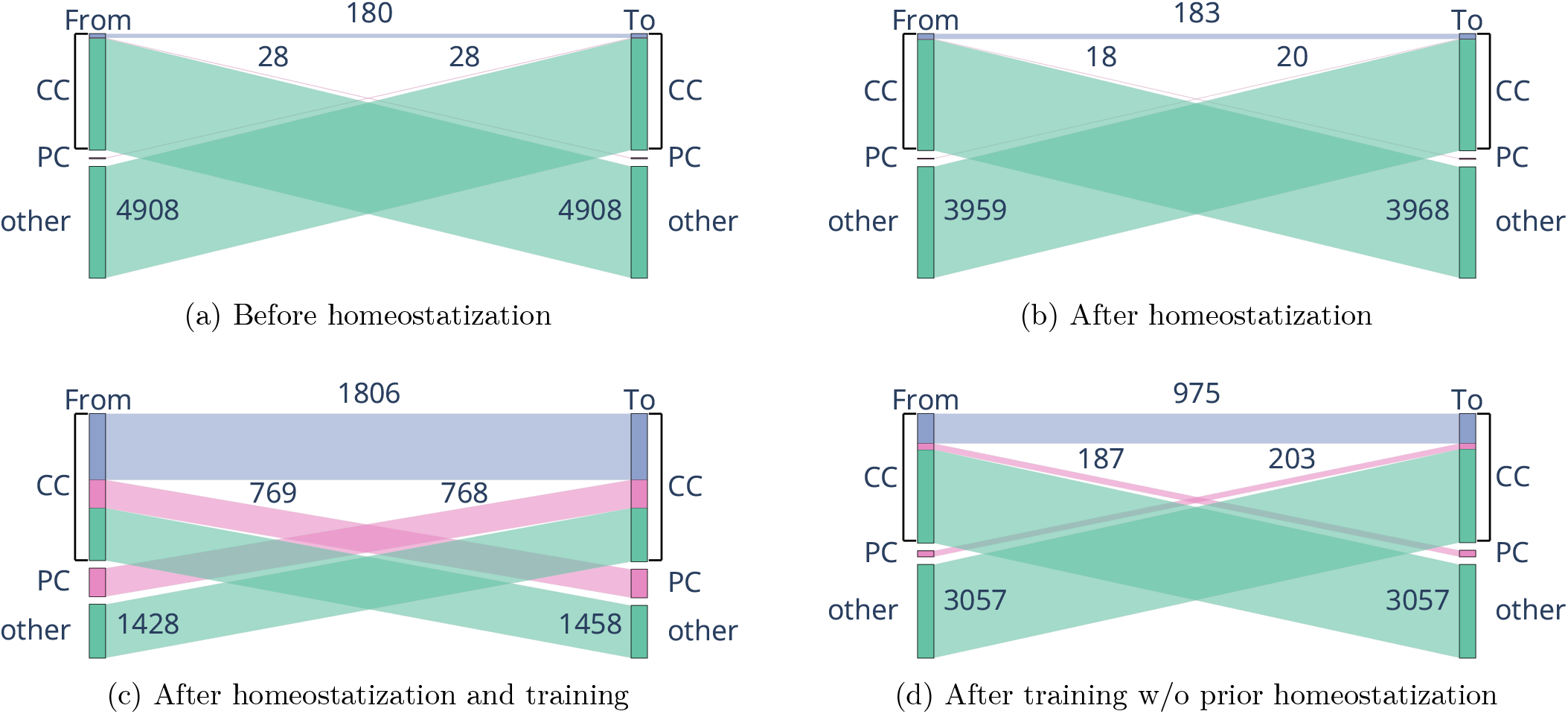
Changes in the number of incoming and outgoing synapses of the CC assembly, including those within it, during homeostatization and training. The rounded numbers represent the average across all 36 connectomes. The subfigures depict the state before homeostatization (a), after homeostatization (b), after training (c), and after training but without prior homeostatization (d). Homeostatization led to a slight net reduction of synapses. Training strengthened connectivity within the CC and between the CC and the PC while reducing connectivity to other regions. Omitting homeostatization substantially diminished the effects of training.

**Fig. 6:**
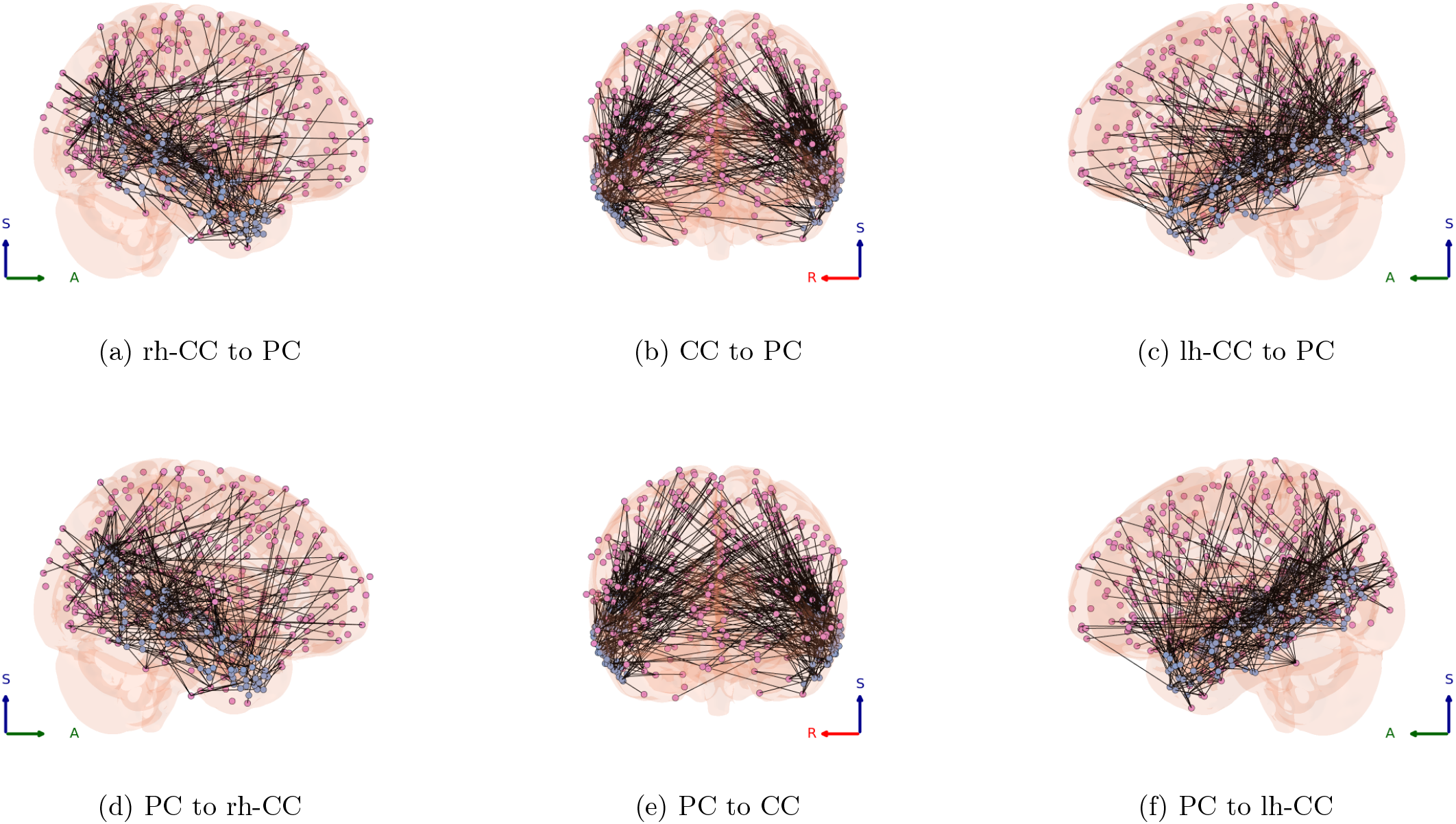
A structural engram after training, viewed from different angles, as indicated by the Right-Anterior-Superior coordinate system in the corner of each subfigure. Newly formed synaptic connections are represented by straight black lines. The PC neurons belonging to the engram are marked in pink and the CC neurons in blue. The prefixes *lh-* and *rh-* stand for left and right hemisphere, respectively.

### D. Recall

To assess the functional consequences of these additional synapses, we examined whether they altered neuronal firing patterns. Specifically, we tested whether CC neurons had learned to respond to PC activation, simulating the recognition of concepts after a virtual sensory experience modeled by manual PC activation. Conversely, we assessed whether CC activation could elicit firing in the PC, thereby mimicking the recall of perceptual experiences associated with a learned concept. This allowed us to identify the *functional engram*, defined as a subset of structural-engram neurons (i.e., neurons at either end of the synapses forming the engram) that respond with high statistical significance to stimulation of the opposite group. The protocol and further details on the calculations are provided in the appendix (Table A3b).

We used the z-score to determine whether a neuron reacted to a stimulus. First, we calculated each neuron’s base frequency. Then, we took the neuron’s firing rate during the stimulation (*f*_*n*_). Finally, we computed how many standard deviations the stimulated activity is away from the mean activity prior to stimulation:

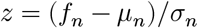

If *z* ≥ 5 or *z* ≥ 11, we concluded that the neuron’s reaction was statistically significant, indicating this using the labels 5σ or 11σ, respectively.

Fig. 7 illustrates the functional response in one connectome, while Fig. 8 breaks this response down by brain area. Moreover, to confirm the necessity of prior homeostatization at the functional level as well, we conducted an ablation study without homeostatization. Again, its absence impeded training. Fig. A3 shows the distribution of results for the 36 individual connectomes, first with a significance threshold of σ = 11. The response signature of the original connectome, before homeostatization and training, serves as the baseline. Clearly, forgoing prior homeostasis resulted in a weak response because an imbalanced connectome generates its own signal, which interferes with the training signal. Lowering the threshold to σ = 5 did not change this pattern; skipping homeostatization not only weakened CC-PC connectivity but also increased spurious responses in unrelated neurons.

**Fig. 7:**
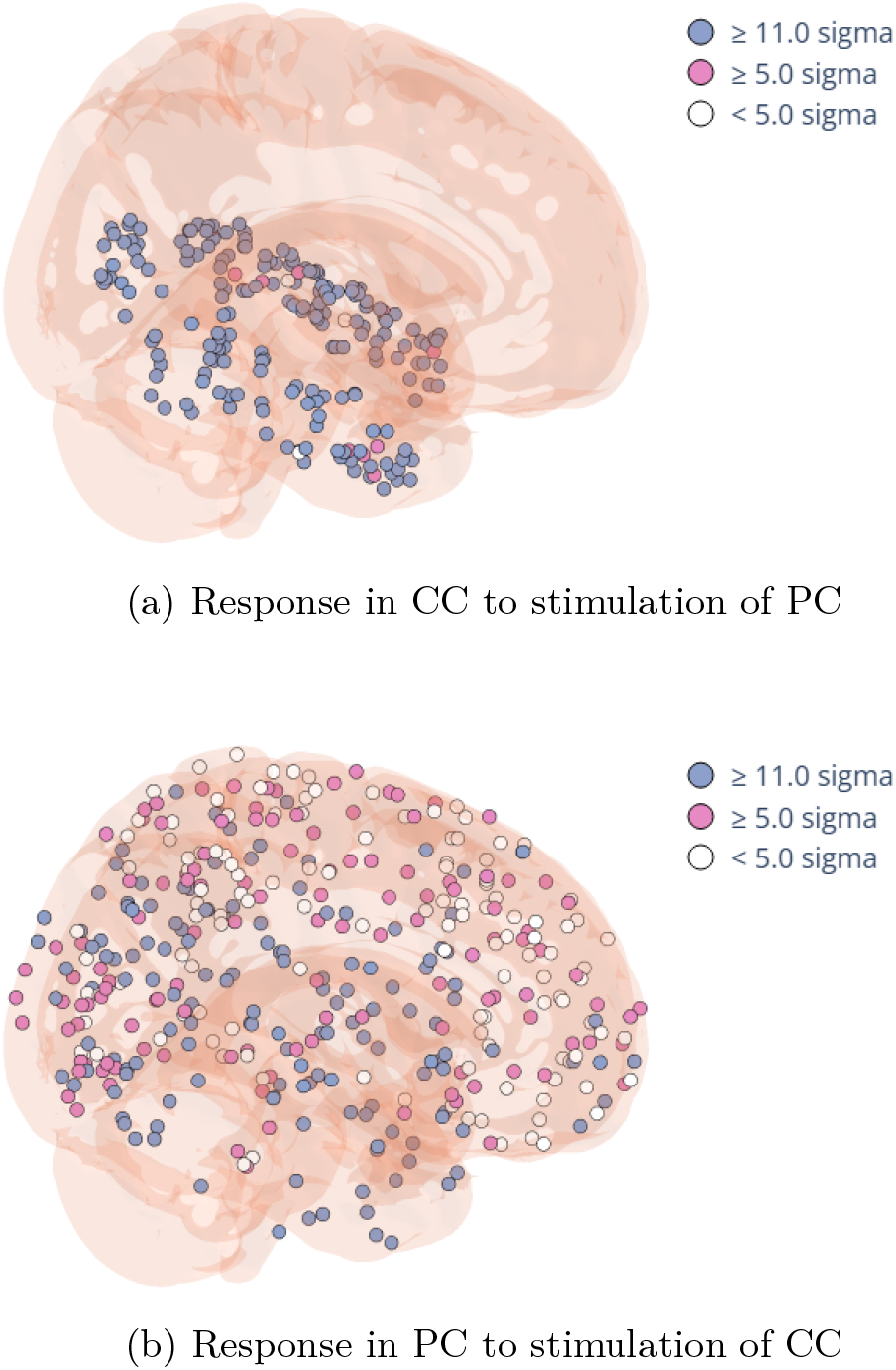
Functional engram after homeostatization and training. The colors indicate the significance threshold a neuron’s response reaches in one assembly when the other is stimulated.

**Fig. 8:**
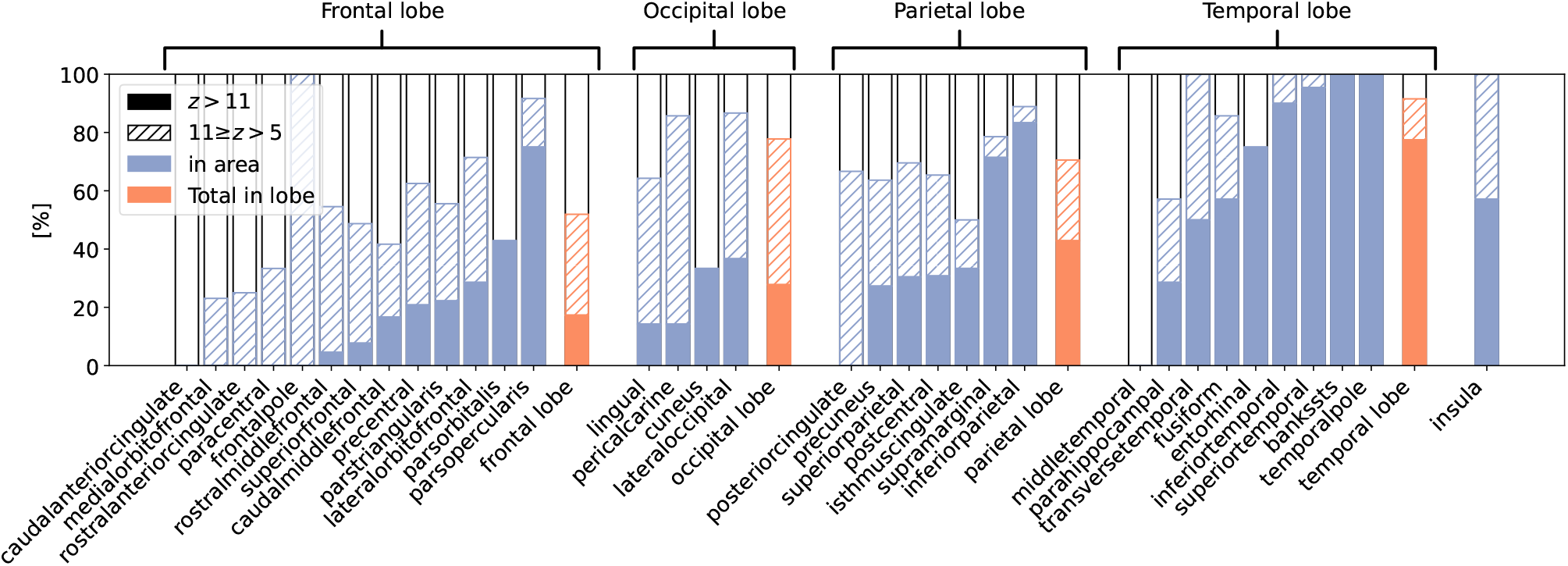
Response of PC to a stimulation of CC, broken down by brain area according to the Desikan–Killiany atlas and ordered in ascending order by the relative number of responding neurons. We consider the sum across both hemispheres. Solid bars mark the response with high significance of *z* ≥ 11σ, and hatched bars mark the medium range of *z* ∈ [5σ, 11σ). Blue bars indicate the response in individual brain areas, while orange bars indicate the response in an entire lobe.

### E. Multiple engrams

Since the human brain can store multiple concepts, we expanded our study to include more than one engram per connectome. In our structural connectomes, only about 1500 out of 47 000 neurons belong to the MTL, limiting the number of concepts we can train. This is especially true since not all MTL neurons should be allocated to concepts. We modeled multiple engrams as follows: Concept cells belonging to different engrams formed disjoint subsets, meaning each MTL neuron belonged to at most one concept, whereas percept cells were allowed to overlap across different engrams, however unlikely such an overlap may be. We trained five engrams, again, each with 200 concept cells and 400 percept cells. We used the protocol in Table A3a, instantiated for five engrams. In the appendix (Fig. A1), we show how the connectomes changed the number of their synapses during training. Again, synaptic shedding preceded the formation of new synapses. We also evaluated the functional response after training (Fig. A2). Stimulating the PC assembly within the same engram produced a strong response in the corresponding CC assembly. Compared to experiments that trained only a single engram, the reverse functional connection from a CC to its corresponding PC was weaker yet still discernible. As expected, the response in the neuronal population outside of any engram was relatively low. However, when training a larger number of engrams, it is impossible to isolate the effect of explicit homeostatization, as we did in the single-engram scenario. Stimulating all the neuron assemblies during the first phase of the protocol—prior to their pairwise stimulation—takes so long that it implicitly homeostatizes the connectomes anyway.

### F. Percept-concept loops

Finally, we applied our framework to simulate percept-concept loops, a virtual thought process designed to model free association—the spontaneous and unfiltered chains of thought common to human mental experience. The loop begins with a set of perceptual features derived from current sensory input. Once stimulated, those features may trigger one or more concepts, which, in turn, may trigger other mental images or memories. Based on those, the subject may again be reminded of further concepts, and so forth. The result is a mental loop oscillating between two different levels of abstraction, slightly varying the percepts and concepts touched in each iteration until it is interrupted by a dominant input that absorbs all attention. Free association is a well-known method in psychoanalysis, used to uncover hidden (i.e., unconscious) memories. In our model, these hidden memories appear as latent percept-concept links to be discovered in such a loop. Let us consider an example. Looking at a family photo (Percept 1) reminds us of grandma (Concept 1). This evokes memories of her cookies, their taste, and texture (Percept 2). These memories trigger thoughts about the winter season (Concept 2), a time when we enjoyed these cookies. In this case, grandma’s cookies serve as a shared percept that connects the two concepts, *grandma* and *winter season*, each with its own associated perceptions. Although we let the loop start here with a percept, as a trigger to activate the first concept, such a loop could equally begin with a concept. What matters is the percept-mediated transition between concepts.

In our simulation experiment, we used a single home-ostatized and trained connectome containing two concepts. Each concept consisted of 200 concept neurons and 400 percept neurons. The two percepts overlapped by one-third, meaning they shared 133 neurons, while 267 percept neurons were unique to each concept engram. The concept neurons of the two engrams, however, were completely disjoint. In later tests, we varied the overlap ratio between the percept assemblies of the two engrams.

Fig. 9 illustrates the transition from one concept to another with an overlapping set of perceptual features (PC1 → CC1 → PC2 → CC2, PC1 ∩ PC2 ≠ ∅). The cell assemblies are enumerated depending on the concept they belong to. In the first step, we stimulated the perceptual features related to the first concept but not those related to the second (PC1 \ PC2). This activated a significant number of concept neurons in CC1, with over 90 % reaching the 5σ significance threshold. Next, we stimulated this exact portion of CC1, eliciting a significant response in a subset of PC2 neurons, particularly those at the intersection of the two feature sets (42 % of the 133 neurons shared between PC2 and PC1 but only 13 % of the 267 neurons unique to PC2). Subsequently, stimulating this subset evoked thoughts of the second concept (CC2), activating a substantial proportion of its concept neurons (75 %). Recalling grandma (CC1) after activating some but not all sensory features related to her (PC1 \ PC2) brings to mind the smell of her cookies, which lies at the intersection of PC1 and PC2 but was not present when the association chain began. The cookie smell then triggers the concept of the winter season (CC2), possibly evoking further memories, such as snow and candles (PC2 \ PC1).

**Fig. 9:**
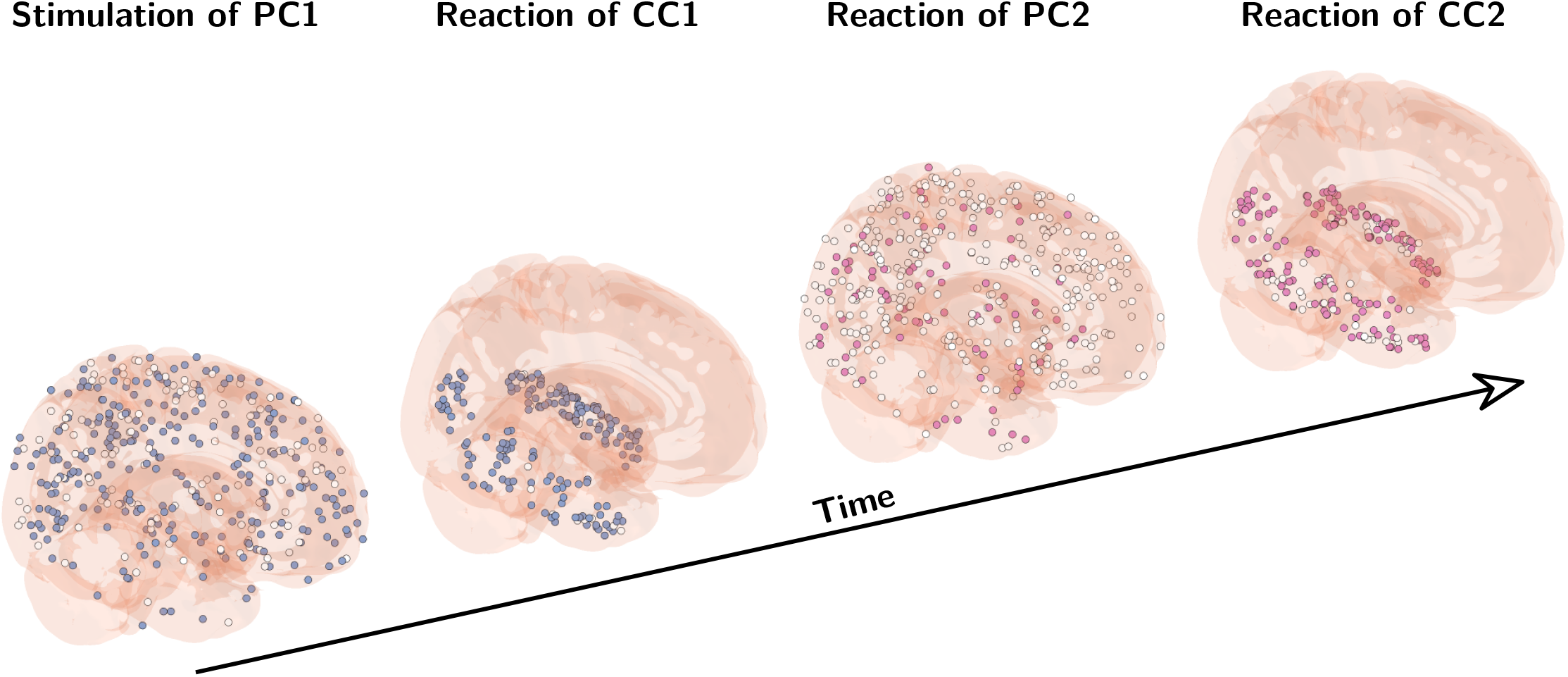
A percept-concept loop involving two engrams, (CC1, PC1) and (CC2, PC2). The loop begins with the stimulation of PC1, excluding the percept neurons that also belong to PC2. Those neurons in each group that react with at least 5σ are colored, while those that do not react significantly are left white. Blue neurons belong to the first engram, while pink neurons belong to the second.

Following our stimulation protocol, we began by stimulating the first group of neurons with 20 mV for 2000 ms, using the same duration and intensity as in Secs. II-D and II-E. Then, we recorded which and how many neurons responded to this stimulus with significance of at least 5σ and stimulated those that had shown a significant reaction in the next group. We repeated this process until reaching the final group, all the while keeping structural plasticity disabled to prevent changes to the connectome.

The charts in Fig. A4 in the appendix provide a more complete picture of possible transition paths between concepts. They show the quantitative response in the cell assemblies involved when moving from one concept to the next in a percept-concept loop. Fig. A4a shows transitions between different concepts. Fig. A4b shows the oscillation between the same concept and its associated percept for comparison. Since a concept engram trivially overlaps itself to the maximum possible extent, this experiment serves as a baseline for the transition intensity.

Clearly, the transition potential between two concepts depends on the ratio of neurons they share. Table A2 in the appendix shows the results for varying degrees of overlap. Surprisingly, even zero overlap can result in a faint jump between two concepts via unintended cross-talk, reminiscent of having an unexpected thought. We observe that the step from CC1 to PC2 is the weakest link. However, once a critical mass of neurons is reached in PC2, transitioning to CC2 further amplifies the reaction.

## III. Discussion

In this work, we demonstrated how the brain can generate more enduring representations of abstract concepts by rewiring synapses. We also showed how engrams with overlapping percepts interact in percept-concept loops. Attention shifts from a concept to associated memories, which then prompts the activation of another related concept, and so on. For the very first time, the learning process of a personalized brain model, followed by a thought process that leverages the trained capability, has been simulated. Key to this simulation has been the neuronization of the measured connectome, followed by homeostatization to calibrate it for the training task, and the use of a distributed probability kernel for synapse formation. With these tools, we can determine and visualize how abstract concepts are grounded in sensory input. Although our simulation is obviously a coarse approximation, facilitated by connectomes at a fraction of their natural resolution and highly simplified stimulation protocols, it nevertheless retains a subject’s individual traits because our kernel derives changes from existing structures, allowing only minor adjustments to the connectivity. The idea that the interaction between stimulation and response can modify the brain’s structure suggests that computation in the brain occurs on different time scales. At one end of the spectrum is rapid, dynamic information processing, represented by the exchange of individual spikes among neurons. At the other end of the spectrum are structural transformations of the connectome that lead to the gradual stabilization of cognitive functions. Although synapse formation is slower than generating action potentials, it can proceed quickly enough—on timescales of minutes to hours—to keep pace with experience, consistent with the rapid synapse formation observed in motor learning [42]. In a sense, the reverberation of the perturbation forms the engram. Imagine a stone falling into cooling wax: while the wax is still molten, it forms waves that solidify as it cools, moving far more slowly than the stone itself yet still capturing the essence of the event. Similarly, stimulus-induced activity instantly initiates a morphogenetic process that generates a lasting memory trace, even though that process is slower than the perturbation itself.

Our model not only defines a new learning paradigm but also predicts where in the brain such structurally consolidated engrams actually form. Although homeostatic structural plasticity would enable synapse formation between any pair of synchronously stimulated neurons in the brain, the kernel function, derived from the individual subject’s connectome, restricts the possible space. Because we used high-resolution connectomes of healthy human subjects to inform the development of neuronal connectivity, we could predict the brain regions in which newly formed temporal-isocortical engrams would localize. We see the majority of engrams forming between concept cells and neurons in the infratemporal lobe and temporal pole, including the amygdala, which are important relay stations between hierarchically organized isocortical and hippocampal networks [16], [43]. Interestingly, the model predicts engram locations not only within the temporal lobe, but also in the medial orbitofrontal and caudal-anterior cingulate areas. The prefrontal cortex, in particular, seems critical for memory consolidation [43]. Additionally, the model’s engrams may involve higher association networks in sensory cortices, such as the precuneus and supramarginal cortex. This is consistent with experimental evidence on hippocampal-cortical connectivity [44]. Overall, most engrams form in the temporal lobe, followed by the parietal, frontal, and occipital lobes. This makes perfect sense, as concept cells do not receive direct sensory input, but rather respond to highly processed perceptual information from sensory integration sites.

Furthermore, we demonstrated that engrams are not only structural memory traces, but also exhibit stable, reciprocal functional activations of concept cells in the MTL and percept cells in the isocortex. The percept-concept loops we observed, the activation chains of percept and concept cells in the MTL and isocortex, respectively, can be seen as the neural correlate of cognitive association chains involving multiple concepts and percepts. Within the same engram, concept cells responded more fully to activations of percept cells than vice versa. The resulting response of the percept cells was comparatively sparse, consistent with the fragmented sensory memories evoked when merely recalling a concept like grandma.

This directional asymmetry naturally reflects the hierarchical view of abstract representation: concepts in the brain emerge from the further abstraction of highly processed, multimodal perceptual memory traces. Hence, concept cells sit at the top of hierarchically organized circuits that represent not what is perceived, but rather, what is known [16]. In line with previous studies, emerging functional engrams are distributed across multiple brain regions [45], [46]. In our model, the percept cells that define a concept can be interpreted as encoded embedding vectors, which are similar to the mental representation described by Du et al. [47]. This makes it easy to understand how overlap between percepts linked to one concept can trigger a related concept proximate to it in the embedding space. Consequently, association chains are driven primarily by oscillations between concepts and perceptual memories rather than by cells shared between concepts, as Gastaldi et al. suggest in their study [48]. We believe this oscillation more closely reflects the conscious experience of such chains. However, their reasoning regarding the ideal overlap between concepts remains valid; the overlap now appears among percept cells instead. The chain of brain states we observe in our large-scale model resembles conceptual frameworks, such as confabulation theory, in which initial input leads to subsequent brain states arising from “competition” between different symbols [49].

We argue that homeostatic structural plasticity, rather than Hebbian synaptic plasticity, is the more likely mechanistic substrate. In our simulation, concept engrams form in a self-organizing manner in response to associative stimuli. Homeostatic structural plasticity is well suited to forming synapses between previously unconnected neurons in an associative, Hebbian manner, spanning long distances without the need for a global observer of synchronous neuronal activity. Hebb already conjectured that memory formation involves associative strengthening of existing synapses as well as the de novo formation of synaptic contacts [24]. Due to the success of Hebbian learning rules for artificial neuronal networks and methodological limitations in monitoring structural network rewiring in the brain over longer timescales, Hebbian synaptic plasticity has been the predominant subject of study for decades and has become the established doctrine for memory formation and learning. Yet, Hebbian synaptic plasticity has some unsolved shortcomings and would be even physiologically implausible if it were the only plasticity mechanism in the brain. Hebbian plasticity requires an antagonist to prevent runaway excitation, typically achieved through homeostatic plasticity. Hebbian plasticity performs well in fully connected artificial neural networks, but it can lead to suboptimal performance in sparsely connected networks, particularly in changing environments. Even worse, networks with pure Hebbian synaptic plasticity are at risk of catastrophic forgetting [50] when they exceed their memory capacity [51]. Homeostatic structural plasticity seems to be the missing puzzle piece that accounts for engram formation and elegantly circumvents the unresolved issues of traditional forms of Hebbian synaptic plasticity. In fact, the enlargement of existing synapses induced by LTP after theta-burst stimulation is accompanied by the pruning of dendritic spines in the vicinity of amplified synapses, implying a homeostatic regulation of neuronal activities by structural processes [52].

Our hypothesis is bolstered by additional empirical evidence and logical reasoning. First, de novo synapse formation has been observed in living organisms, even over longer distances [7]. De novo formation of synapses is required in sparsely connected networks when novel associations are formed, typically when novel concepts emerge from the integration and abstraction of highly multimodal and multicategorical perceptions. Essentially, concept cells in the MTL must quickly form cell assemblies with remote neurons distributed across various sensory isocortical association areas after only a few presentations. However, the current Hebbian doctrine cannot explain how such remote cell assemblies form in a goal-directed manner. Goal-directed Hebbian formation of de novo synapses would require all-to-all connectivity or a global observer of electrical activity—an implausible assumption. Changeux and Danchin postulated in their selective stabilization hypothesis that synapses form randomly during development and are either consolidated by Hebbian synaptic plasticity or pruned later [53]. However, all later variants of this notion fail to explain how new connections can form specifically in response to the synchronous stimulation of unconnected, remote neurons. In contrast, homeostatic structural plasticity can form new synapses in a Hebbian manner and, as demonstrated here, can serve as a goal-directed mechanism for associative synapse formation over any distance in the brain, provided that some precursor connections are available. By adding synapses and pruning interfering synapses, it may quickly generate a large, orchestrated impact on downstream neurons, so that even single-shot learning seems possible through homeostatic structural plasticity. Indeed, the rapid formation and selective stabilization of new spines has been demonstrated in the motor cortex when acquiring novel motor skills, even in adulthood [42]. Second, both operational and computational efficiency demand structural plasticity. Adapting connectivity to tasks by introducing shortcuts where needed while abandoning inactive circuits increases processing speed without incurring additional energy or material costs. Finally, this solution is more stable, enabling the instant recovery of skills learned during childhood, such as riding a bicycle or skiing, even after long periods without practice. Beyond mere plausibility, our mechanistic view also yields specific, testable predictions about how engrams form in the living brain. Although the formation of structural elements and the consolidation of synapses are intertwined, the sequence of events can be distinguished by the type of plasticity—Hebbian synaptic or homeostatic structural—that drives engram formation. With Hebbian synaptic plasticity, one would expect increases in dendritic spine and axonal bouton densities, as well as connectivity, that scale with the extent of stimulation. These effects are counterbalanced by subsequent homeostatic scaling of synaptic strengths and pruning of connections. With homeostatic structural plasticity, however, we would expect to observe the opposite. Stimulation initially triggers the pruning of connections, followed by the formation of axonal boutons and dendritic spines, as well as an increase in connectivity. Pruning and increased connectivity would scale with stimulation strength and duration; and yet, activities would not exceed homeostatic regimes because the entire process is homeostatic in nature. The two plastic processes have different characteristics and should be distinguishable in the living brain.

Yet, empirically validating our model of homeostatic structural plasticity beyond the mere sequence of pruning and growth is non-trivial. Tracking individual synapses in vivo poses immense technical and ethical challenges, especially in humans. Furthermore, we must translate complex sensory inputs into discrete digital activations while accounting for the avatar’s lower structural resolution. Ultimately, validation requires a statistical homomorphism—a mapping that preserves the brain’s functional algebra in its avatar. This would allow us to generate equivalent inputs and reasonably compare outputs, thereby testing whether the avatar’s distribution of structural and functional responses mirrors biological reality.

The dynamic interplay between perception and concept is also essential to understanding artwork. In poetry, verses evoke concepts that stimulate mental images or memories, which then stimulate further reflections in our semantic-perceptual network. A *leitmotif* [54] is a recurring musical theme in an opera that represents a character, person, idea, or situation. By closely linking a concept to sensory signals, a leitmotif can construct powerful, multimodal scenes of striking emotional intensity. Generally, the semantic properties of an object—a piece of art, in this context—are determined by its relationship to a potentially infinite array of other objects [55], whether conceptual or perceptual in nature. Our model describes the iterative exploration of these semantics following a chain of memories and mental symbols. While we do not assert that every exposure to artwork fosters the formation of new synapses, our model illustrates how such perception could occur based on concepts trained earlier in a different context. Nevertheless, it is not inconceivable that, after frequent or intensive exposure, a particular sound or image may eventually provoke a hardwired response.

Our study offers a novel perspective on the role of neural network structure in supporting learning processes. In our simulation, we injected this process into the connectome of an avatar representing a real person. In a sense, our work reverses the conventional concept of a brain-computer interface. The computer controls a virtual copy of a person’s brain. While this has been previously explored in different ways [56], [57], [58], demonstrating the learning of concepts based on structural plasticity and the recall of learned concepts—including the discovery of related concepts via shared perceptual grounding—has not been accomplished before. The notion of *Gedankenexperiment* (thought experiment) appears in a new light.

## IV. Methods

We simulated the formation of concept engrams in the connectomes of healthy subjects, obtained using the workflow in Fig. 10. Below, we describe the details of image acquisition and tractography, followed by the static and dynamic elements of our brain modeling approach, including efficiency considerations.

**Fig. 10:**
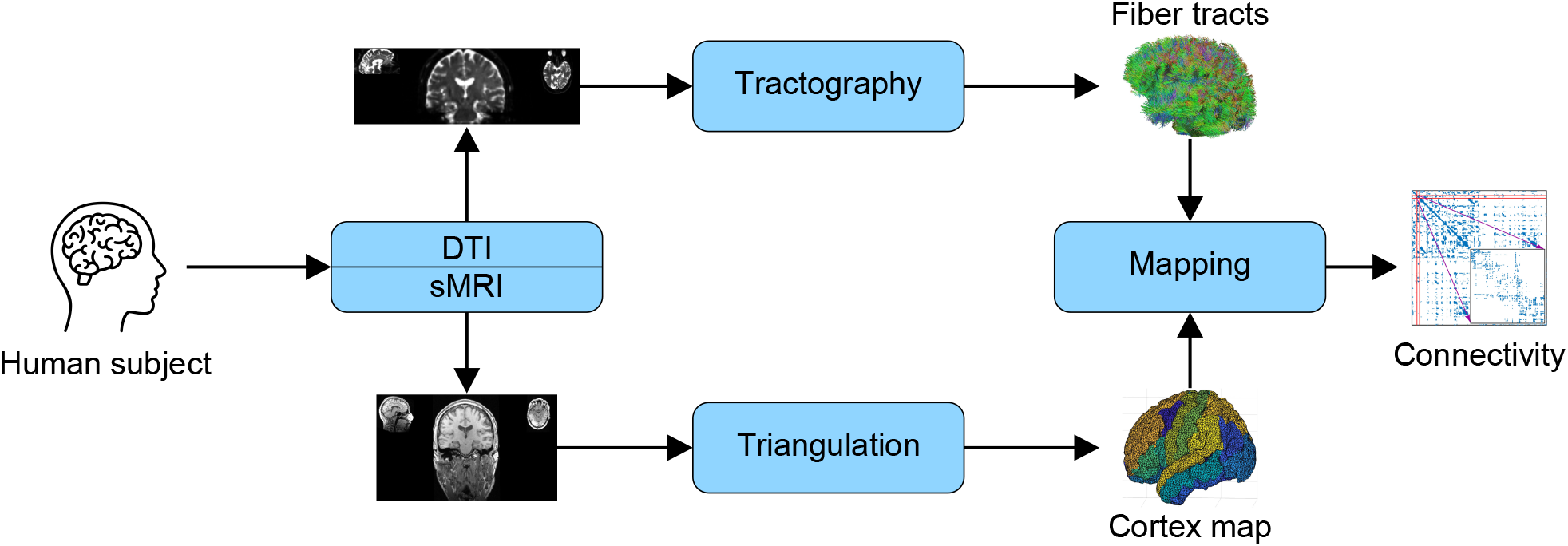
The workflow for obtaining structural connectomes. We used a combination of diffusion tensor imaging (DTI) and structural magnetic resonance imaging (sMRI), followed by high-resolution fiber tractography. To identify inter- and intra-regional connectivity, we parcellated the cortical surface using the Desikan–Killiany atlas [34] to localize brain regions. Furthermore, we triangulated the cortical surface into over 47,000 triangles. We then used these triangles as the seeds and endpoints of streamlines within and between the atlas’s brain regions. Human head on the left: Pavlo S/Shutterstock.com.

### A. Image acquisition

We used structural connectome data from 36 healthy adults who were included as controls in an earlier study by Chen et al. [30]. The connectomes were derived from structural magnetic resonance imaging (sMRI) and diffusion tensor imaging (DTI) originally acquired by Hutchings et al. [59] on a 3T system. For sMRI, they used T1-weighted images with an isotropic voxel size of 1 mm, a repetition time (TR) of 2500 ms, an echo time (TE) of 3.5 ms, and a field of view (FOV) of 256 mm. For DTI, they obtained scans with an isotropic voxel size of 2 mm, a TR of 10 000 ms, a TE of 91 ms, and an FOV of 256 mm with 64 diffusion directions (b = 1000 s mm^−2^) and 12 b0 images. Chen and colleagues used FSL^1^ for standard sMRI and DTI preprocessing of the raw images, including the correction of susceptibility artifacts with reversed-phase encoding directions using FSL’s topup tool^2^.

### B. Tractography

Chen et al. performed whole-brain tractography in *DSI Studio*^3^ using deterministic streamline tractography from preprocessed DTI data. They processed the diffusion data for fiber tracking using a diffusion sampling length ratio of 1.25, then computed the orientation distribution functions on an eight-fold tessellated sphere. This allowed them to resolve up to five fiber orientations (peaks) per voxel. They performed tractography using whole-brain seeding with a quantitative anisotropy threshold of 0.6 × Otsu’s threshold, a turning angle threshold of 60^°^, and a streamline length constraint of 100–300 mm. Of the total 10 000 000 streamlines saved, only those with endpoints in gray matter were retained. To localize the start and endpoints of the streamlines in gray matter, they constructed approximately 47 000 triangles of the brain’s pial surface, each with an average surface area of 3.5 mm^2^. They reconstructed and down-sampled the pial surface from sMRI using an output ratio of 0.1 using the MATLAB toolbox IsoMesh [60].

According to Taylor et al. [61], every triangle of the surface represents one node of the high-resolution structural connectome. The structural connection strength between two nodes was measured by the normalized streamline counts running from one corresponding triangle to a second triangle. Based on the nodes’ locations on the pial surface, we assigned each node of the high-resolution structural connectome to one brain region of the Desikan– Killiany cortical parcellation [34]. With this assignment, each cortical area contains hundreds of nodes and can form an intra-area network.

### C. Brain model

While the previous study has assessed the clinical relevance of brain connectivity for epilepsy, our goal is to establish normative intra- and inter-regional connectivity patterns in healthy brains to inform the layout of the avatar connectomes for our engram study. Each of our avatar connectomes consists of approximately 47 000 spiking neurons. Their positions in 3D space were derived from the center-of-mass coordinates of the triangulated cortex. We decided to map the center of mass to neurons one-to-one to avoid introducing side effects when choosing neuronal densities across cortical areas. However, this means that each neuron in our avatar represents a multitude of neurons in the actual cortex. The exact number of neurons per avatar brain varies across subjects. Neuronal activity follows the Izhikevich model [62], which provides a good trade-off between realistic firing patterns and computational effort. Initial network connectivity was derived directly from each subject’s DTI tractography. Each bidirectional, binary connection from the tractography was converted into two directed synaptic connections in the model, resulting in a symmetrical connectivity matrix with discrete values.

Initially, each synaptic connection had a weight of one. In general, our model allows multiple synapses per neuron pair, corresponding to weights greater than one, an option present in our simulations. Additionally, we allow the connectivity matrix to be asymmetrical, meaning the number of synapses between two neurons in one direction is independent of the number in the other direction.

In our model, connectivity is not static. It changes over time in response to activity patterns, following the model of structural plasticity by Butz and van Ooyen [12], which we describe in more detail below.

### D. Model of structural plasticity

The model of structural plasticity (MSP) simulates a reciprocal interaction between activity-dependent adaptations of neuronal morphology and the rewiring of connectivity, thereby altering neuronal activity over longer timescales. MSP simplifies neuronal morphology by representing contact interfaces, such as axonal boutons and terminals, as well as dendritic spines and postsynaptic receptor densities, as axonal and dendritic elements, respectively. These, collectively called synaptic elements, develop independently and increase or decrease in number in an activity-dependent fashion. Vacant axonal and dendritic elements offered to the neuropil are matched to form new synapses. Matchmaking is random, based on the supply and demand of synaptic elements, and favors neurons that are close to one another. For instance, neurons with a high number of vacant dendritic elements are likely to be targeted by nearby axons in search of a new partner. However, as we will explain in Section IV-F, we modified this rule to create a more realistic model of how new long-range connections form. Losing an axonal or dendritic element bound in a synapse immediately causes the synapse to break. MSP enforces a reciprocal interaction between neuronal activity and connectivity. Imbalances in postsynaptic firing rates trigger morphological adaptations that increase or decrease the number of axonal and dendritic elements. These changes in neuronal morphology then cause rewiring of neuronal connectivity, which may ultimately result in homeostatically balanced neuronal activities. We can draw an analogy with particle physics. Vacant synaptic elements in MSP create a force field that directs axon growth, serving as a mechanism for estimating gradients [63] toward homeostatic equilibrium. MSP was used for both homeostatization and training— with the simulation parameters listed in Table A1.

### E. Growth of synaptic elements

According to the original formulation of MSP, the intracellular calcium concentration [*Ca*^2+^] indicates a floating average of neuronal firing rates. A sigmoidal growth rule determines the growth of synaptic (i.e., axonal and dendritic) elements (*dA, dD*)/*dt*, modeling the growth and retraction of axonal boutons and dendritic spines as a function of a neuron’s average firing rate, abstracting away from detailed neuronal morphologies (Eq. 1 and Fig. 11).

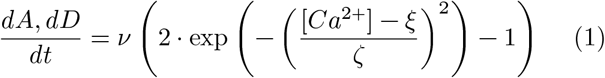

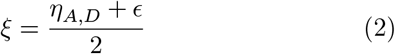

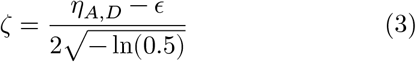

**Fig. 11:**
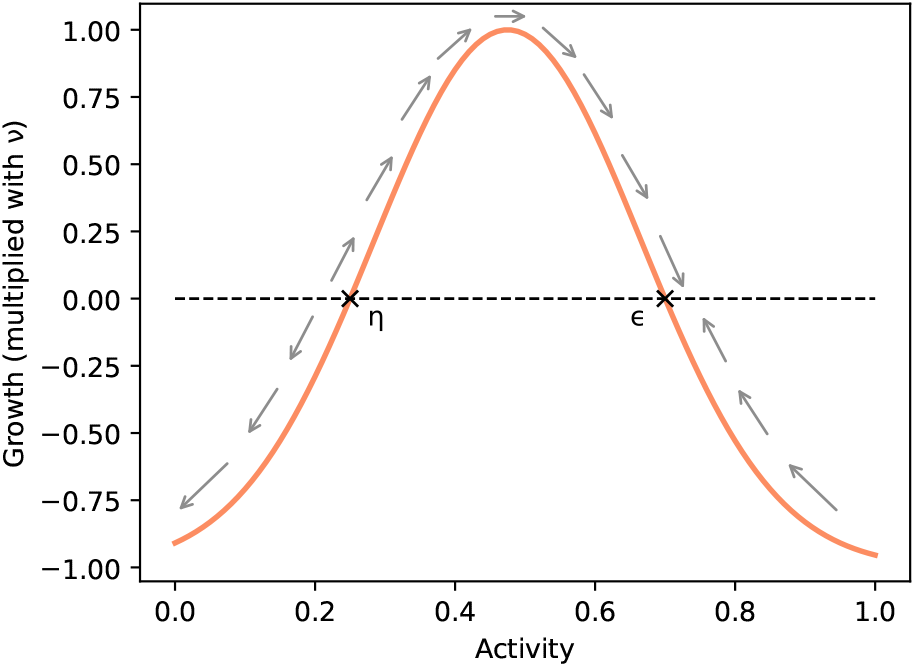
Activity-dependent growth of synaptic elements is described by a Gaussian growth curve. Intracellular calcium ([*Ca*^2+^]) is given as a unit-less concentration on the x-axis and used as a floating average for neuronal firing rates and, hence, the neuron’s activity. The growth or retraction of elements is expressed in fractions of the growth rate *ν*. The growth curve is bound to the interval (−1, 1]. The intersections of the curve with the x-axis are the only fixed points of the system. η = 0.25 is an unstable fixed point and ϵ = 0.7 is a stable fixed point. If a neuron can fulfill its need to form synapses with available synaptic partners, it will develop its activity towards the stable fixed point (gray arrows), which then functions as a homeostatic set-point for the neuron’s electrical activity.

With a growth rate of *ν* = 10^−4^, the fast time scale of activity fluctuations can be separated from the slow time scale of synaptic growth. For each neuron, a stable fixed point was chosen at ϵ = 0.7. All neurons can develop their activities toward this fixed point as long as their initial activity is greater than the unstable fixed point η_*A*_ = η_*D*_ = 0.25 on the left of the growth curve in Fig. 11. Neurons transmit synaptic activation of 1 mV to downstream neurons whenever they fire. In addition, neurons receive background noise modeled as a Poisson process with *N*(*µ*, σ^2^) and *µ* = 5 and σ = 1.

### F. Long-range synapse formation

The original MSP defined synapse formation as a distance-dependent process, using a 2D-Gaussian kernel *K* (Eq. 4) to predict the likelihood that an axonal element of neuron *i* connects to a dendritic element of neuron *j* as a function of the number of vacant dendritic elements at neuron *j* (vac_*j*_) and the Euclidean distance between their host neurons.

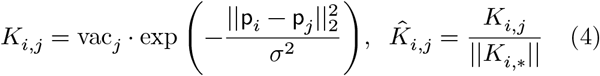

While this approach worked well for local connectivity [15], it altered the network topology of our connectomes beyond recognition. Moreover, the Gaussian kernel makes it rather unlikely that new synapses form over long distances, for example, between neurons in remote brain areas.

Although axons will not grow out farther than a few hundred micrometers in the adult brain [7], [64], de novo synapse formation between remote neurons can indeed occur in the mature brain if axonal terminal branches sprout out or form en-passant synapses with dendritic spines in reach to connect previously unconnected neurons [9]. Depending on axon length, the cell bodies of pre- and postsynaptic neurons can be located in different brain regions.

We extended MSP to enable the formation of long-range connections by introducing a distributed kernel *H* (Eq. 5), which reflects the anatomical layout of projection fibers formed during juvenile brain development. As the anatomical layout of connections is less likely to change in response to learning in an adult brain, we keep *H*’s initial value at *t*_0_ constant throughout the simulation. To initialize *H*_*i,j*_, we center a Gaussian kernel *K*_*i,j*_ (Eq. 4) per neuron at the neuron’s position p_*i*_ as in the original MSP but now in 3D space. The original formulation used 2D positions of neurons in the primary visual cortex. Additionally, we place a further Gaussian kernel centered at the position p_*j*_ of each downstream neuron *j*, to which neuron *i* initially projects to (i.e., for which the initial connectivity 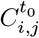 is greater zero (Fig. 12)). We chose σ = 1000 µm for all Gaussian kernels consistently.

**Fig. 12:**
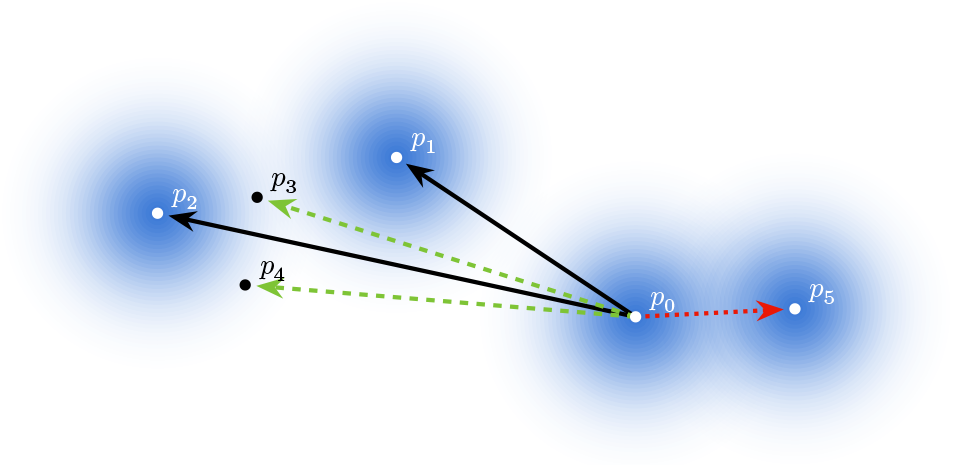
Example for the distributed kernel. In the initial connectivity, neuron *p*_0_ was connected to neurons at positions *p*_1_, *p*_2_, and *p*_5_. In the current step, however, neuron *p*_0_ is connected to neurons at positions *p*_1_, *p*_3_, and *p*_4_. The persistent synapses are shown in solid black. The pruned synapse appears as a red dotted line, and newly formed synapses appear as green dashed lines. Each cloud indicates one summand in the kernel.

During a simulation run, the distributed kernel *H*_*i,j*_ defines the attractivity of a potential target neuron *j* independently for every source neuron *i*. Each potential new target neuron *j*, along with its number of vacant synaptic elements vac_*j*_ in Eq. 4, appears once in every summand (i.e., Gaussian kernel). This is why each kernel evaluates to a value greater than zero for each target.

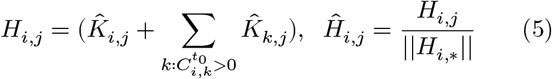

The index *k* ranges over all target neurons, and 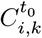 denotes the initial connectivity matrix. With this adaptation (Fig. 12), which we call *MSP with distributed kernel*, the algorithm preserves the measured network topologies in healthy subjects. With Ĥ_*i,j*_, the normalized individual attractivity of every target with respect to the total attraction, we were able to apply an approximation method to reduce the problem complexity of synapse formation in the model, which we explain below.

### G. Efficient approximation of MSP

Matching vacant synaptic elements throughout the network to form synapses is a hard combinatorial problem, the number of possible pairs grows quadratically. With growing neuron numbers and extended simulation times, an efficient approximation of MSP is indispensable. In each update of synaptic connectivity—which occurs every 100 ms—vacant axonal and dendritic elements from neurons *j* and *i*, respectively, form a new synapse with probability Ĥ_*i,j*_. This leads to quadratic complexity if implemented naively. In this study, we used a parallelized approximation of MSP [65], [66]. By exploiting the similarity to N-body problems, the approximation reduces the computational complexity in terms of the number of neurons from *O*(*n*^2^) to *O*(*n* log *n*). This reduction is achieved through an adaptation of the Barnes–Hut algorithm, a widely used method in astrophysics.

Yet, the Barnes–Hut algorithm requires a smooth, rapidly decaying kernel, a condition the composite nature of our novel kernel definition *H*_*i,j*_ violates. However, we could easily address this issue by normalizing each summand in *H*_*i,j*_ to 1, ensuring that each constituent contributes equally to the total. Thus, evaluating the kernel became equivalent to first choosing a summand to evaluate with equal probability and then performing a Barnes–Hut-based search starting at its center, either at the neuron’s own position or at the position of a target neuron to which it had been initially connected. Since each component kernel satisfies the Barnes–Hut requirement, we could proceed as before.

### H. Source code

The source code of our simulator^4^ and the library^5^ we used to calculate the graph metrics is available on GitHub.

## Acknowledgments

Financial support for this work was provided by the following sources: the Federal Ministry of Research, Technology and Space (BMFTR) and the Hessian Ministry of Science and Research, Arts and Culture (HMWK) as part of the NHR funding; the NHR Alliance as part of the NHR Graduate School; the European Union’s Horizon 2020 Framework Program for Research and Innovation (Specific Grant Agreement No. 945539, Human Brain Project— SGA3); the National Natural Science Foundation of China (Grant No. 82301662); and the Natural Science Foundation of Shandong Province (Grant No. ZR2024QH005). The authors gratefully acknowledge the computing time on the Lichtenberg II high-performance computer at TU Darmstadt, which is funded by the BMFTR and the State of Hesse. We thank Michael Reimann for his thoughtful comments.

## APPENDIX

**TABLE A1:**
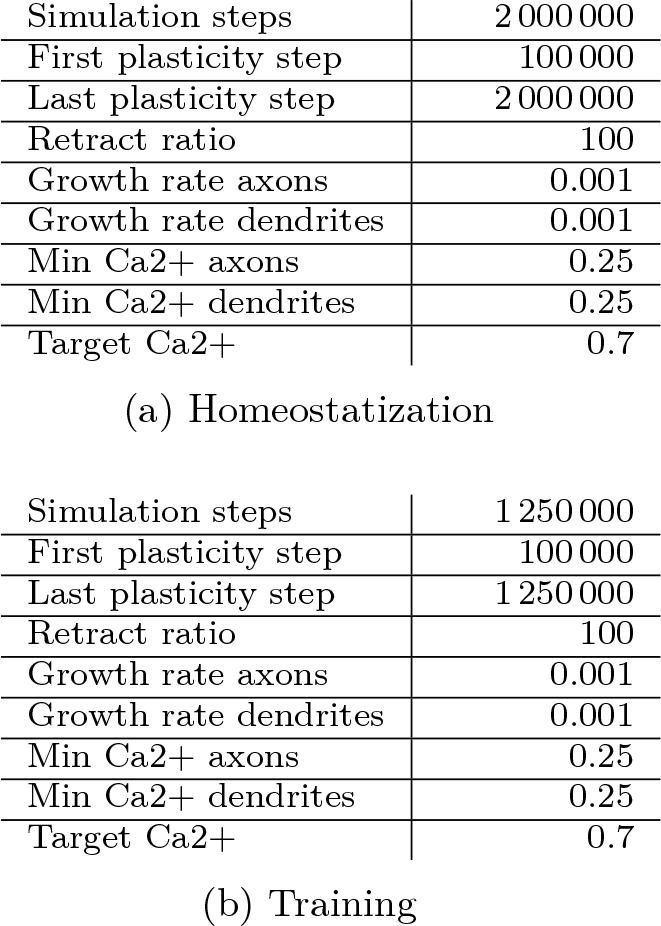
Simulation parameters for homeostatization and training of five engrams. When training fewer than five engrams, we reduced the number of simulation steps in Table A1b by 150 000 for each engram omitted. For details on the timeline of stimulation events, see Table A3a. The first 100 000 steps helped the neurons adapt their connectivity to their electrical activity. On a desktop computer, homeostatization takes approximately 2 h while training takes approximately 1 h.

**TABLE A2:**
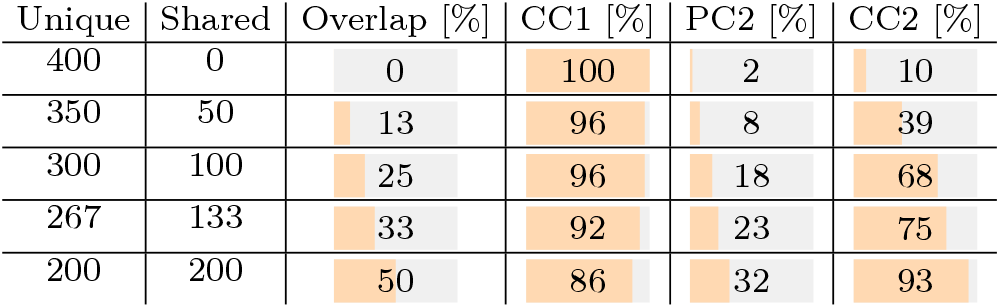
Percept-concept loops involving two engrams with varying fractions of percept neurons shared between them. Every loop started with a stimulation of PC1 \ PC2. The three rightmost columns show the percentage of neurons whose reaction reached the 5*σ* significance threshold as the loop progressed from CC1 and PC2 to CC2. Once the fraction of PC2 neurons activated by CC1 was large enough, transitioning to the second concept became easy.

| Unique | Shared | Overlap [%] | CC1 [%] | PC2 [%] | CC2 [%] |
| --- | --- | --- | --- | --- | --- |
| 400 | 0 | 0 | 100 | 2 | 10 |
| 350 | 50 | 13 | 96 | 8 | 39 |
| 300 | 100 | 25 | 96 | 18 | 68 |
| 267 | 133 | 33 | 92 | 23 | 75 |
| 200 | 200 | 50 | 86 | 32 | 93 |

**Fig. A1:**
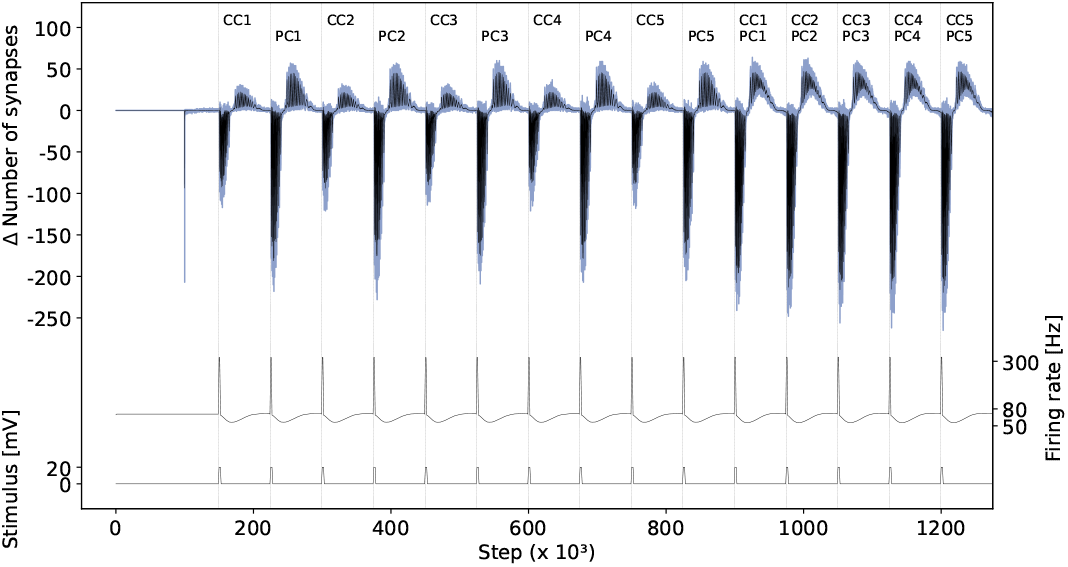
Change of neural activity and number of synapses when training five concepts. According to our stimulation protocol in Table A3a, this involves fifteen distinct stimulation events (bottom). The labels at the very top indicate the assemblies being stimulated in each step. Each event causes the neural activity (i.e., firing rate) in the stimulated assemblies to peak, followed by a wider but shallower slump (middle). In response to activity exceeding the set-point, neurons shed existing synapses to compensate for their increased activity (top). During the subsequent period of low activity, neurons that had previously lost synapses formed new ones until the original activity level was restored. Because all of the co-stimulated neurons were seeking new partners at the same time, they tended to connect with each other. This resulted in the formation of the engrams. The blue region represents the spread of the difference in the number of synapses relative to the previous time step across all 36 connectomes, while the black line represents the average. This number reflects changes throughout the entire connectome, including neurons outside of the stimulated assemblies. Since the stimulation protocol was identical across connectomes, and the networks were nearly identical in size with the same number of stimulated neurons, the traces from the individual connectomes overlap almost completely.

**Fig. A2:**
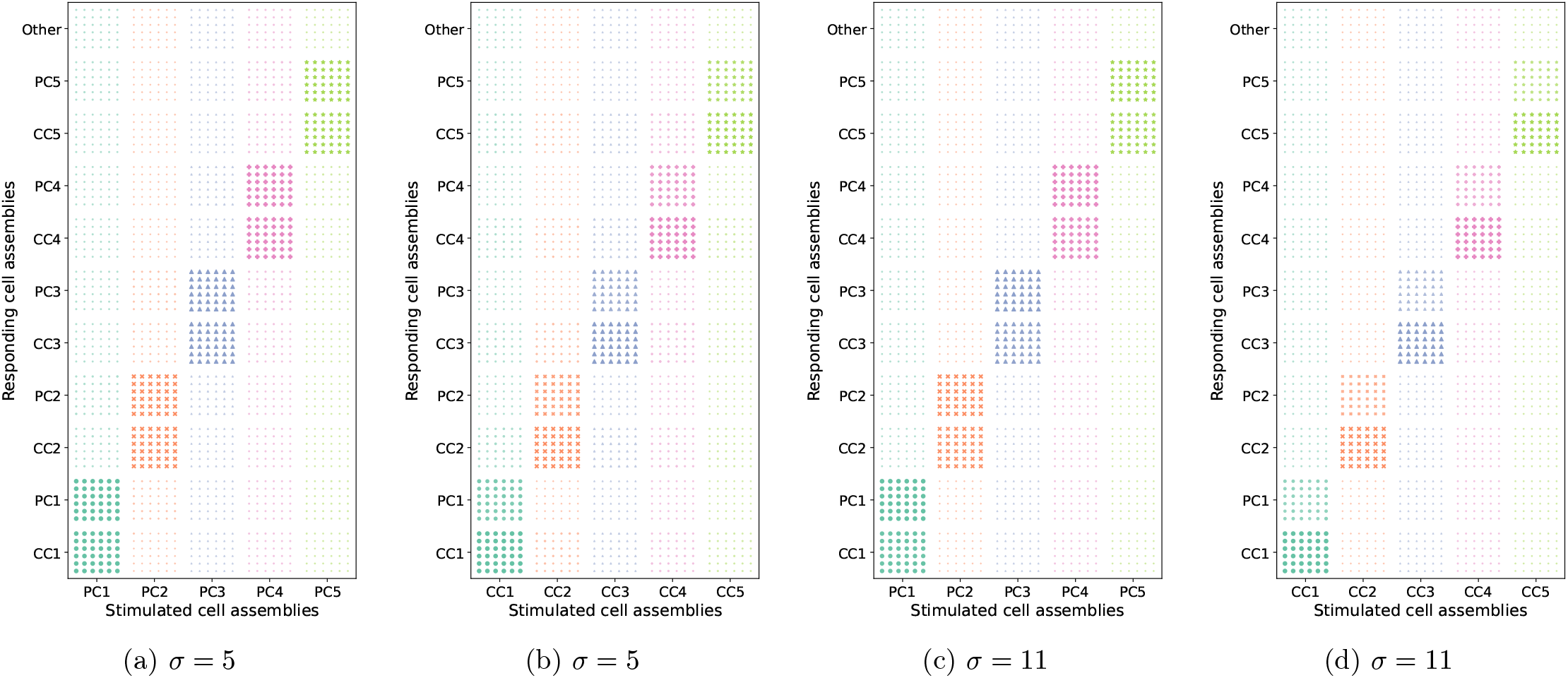
Functional evaluation of five engrams after homeostatization and training across all 36 connectomes. The x-axis shows the stimulated assemblies, and the y-axis shows how the other assemblies respond. For each combination, we plot the reaction intensity of each of the 36 connectomes. This is expressed as the percentage of neurons whose activity reached the significance thresholds σ = 5 and σ = 11, respectively. The percentage is visualized in two ways: the size of the dots and their transparency. The results across all subjects were fairly consistent.

**TABLE A3:**
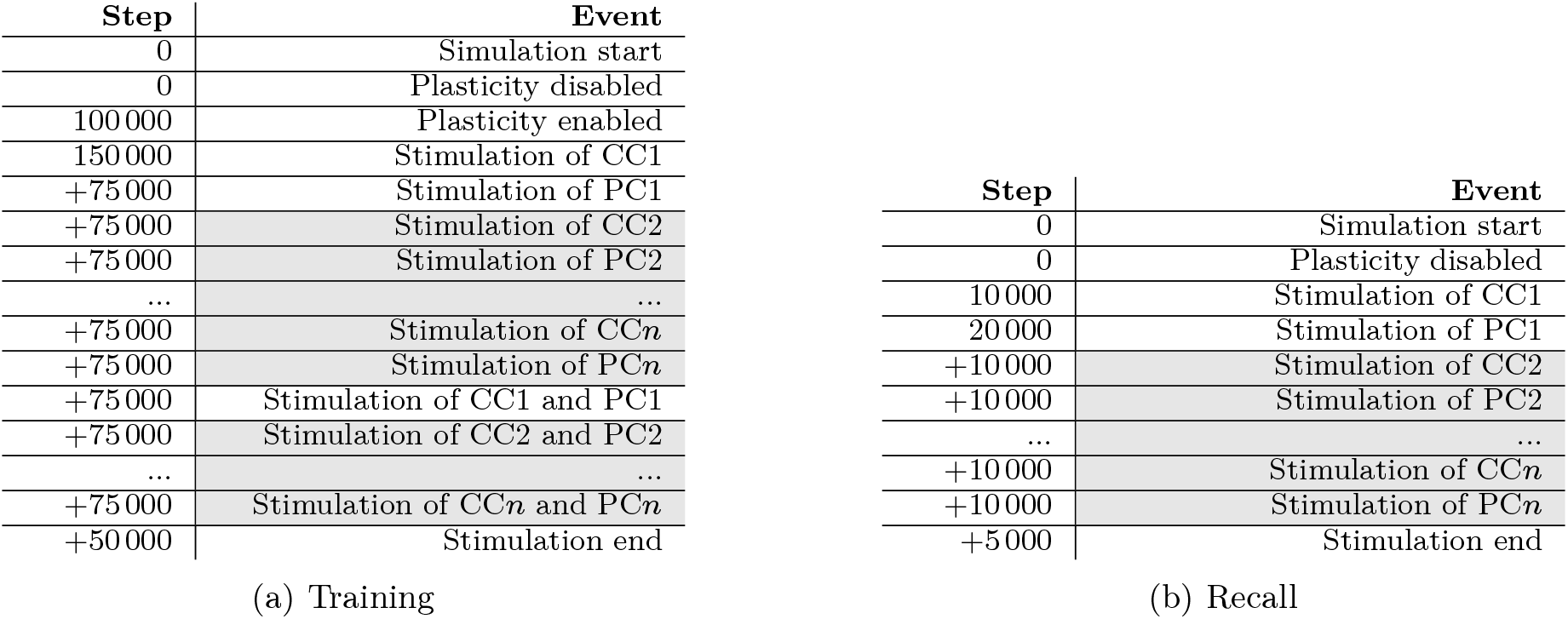
Protocols for training and recall for both single and multiple engrams. The cell assemblies are enumerated depending on the engram they belong to (CC*i* and PC*i* belong to the *i*th engram). The single-engram version is obtained by omitting the gray-highlighted steps. Each stimulation applied 20 mV for 2000 simulation steps. Spacing the stimulation events 75 000 steps apart gave the neurons enough time to adjust their connectivity. During recall, 10 000 steps between stimulation events ensured that individual stimulations did not interfere. We derived the significance threshold in our recall experiments from the baseline neuronal activity (i.e., firing frequency). This baseline is the average frequency between steps 1000 and 9999, calculated separately for each 1000-step interval. Splitting into smaller intervals was necessary to compute the standard deviation.

**Fig. A3:**
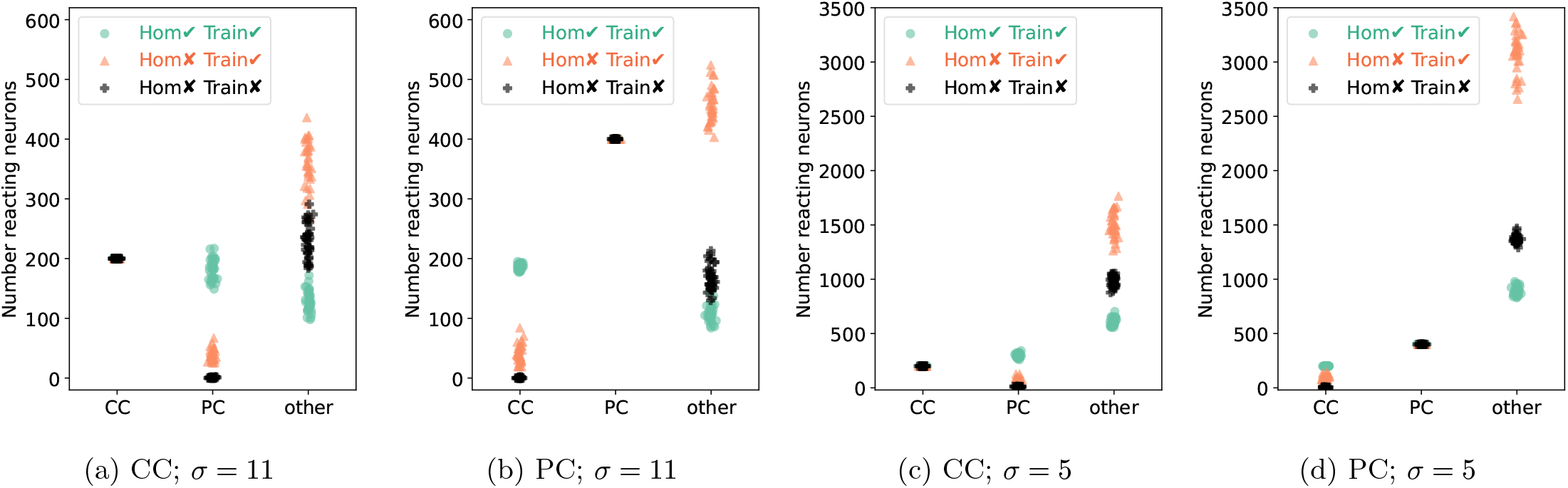
Functional evaluation of single engrams across all 36 connectomes, including ablation results that isolate the effect of homeostatization. The plots show how the assemblies CC and PC responded to stimulation of either CC or PC, as indicated in the subcaption. The data points represent the number of neurons whose responses reached significance thresholds of 5*σ* or 11*σ*. Results are shown after homeostatization and training (green circles), after training but without prior homeostatization (orange triangles), and with neither homeostatization nor training (black crosses). The last-mentioned configuration is the response of the original connectome right after neuronization and serves as a baseline. To reduce overlap, we slightly varied the symbols’ horizontal positions.

**Fig. A4:**
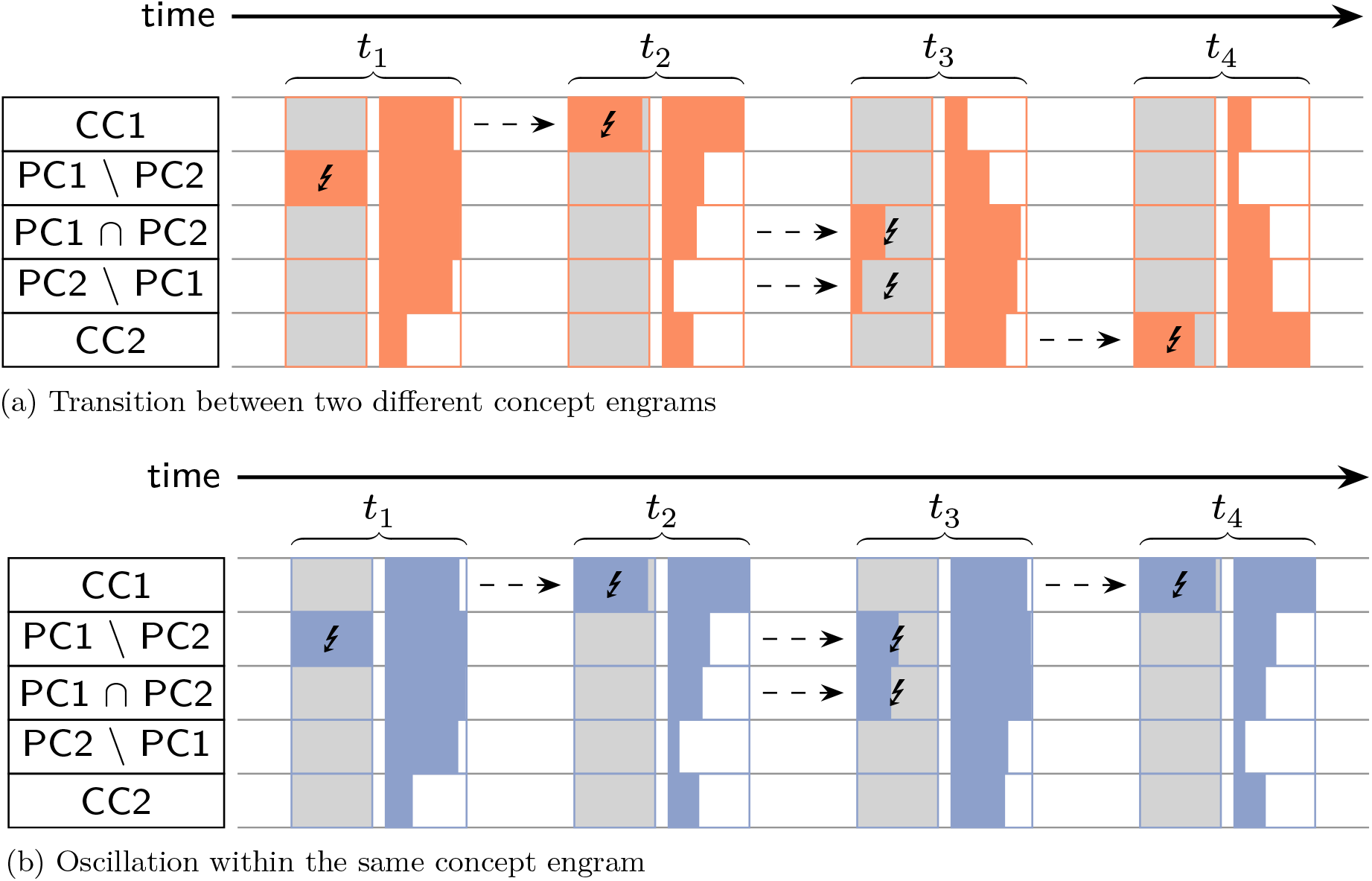
Quantitative analysis of the transition paths between concept engrams. Using a homeostatized connectome with two concept engrams consisting of 400 percept neurons and 200 concept neurons each, we investigated the transition between two different concepts (PC1 → CC1 → PC2 → CC2) in Fig. A4a (orange) and the oscillation within the same concept (PC1 → CC1 → PC1 → CC1) in Fig. A4b (blue). The x-axis represents the timeline, and the y-axis represents the various neuron assemblies. Lightning bolts indicate stimulation. The width of each horizontal bar signifies the percentage of neurons whose responses reached the 5*σ* significance threshold. The self-loop experiment establishes the maximum achievable transition intensity between the involved neuron assemblies and serves as a baseline.

**Fig. A5:**
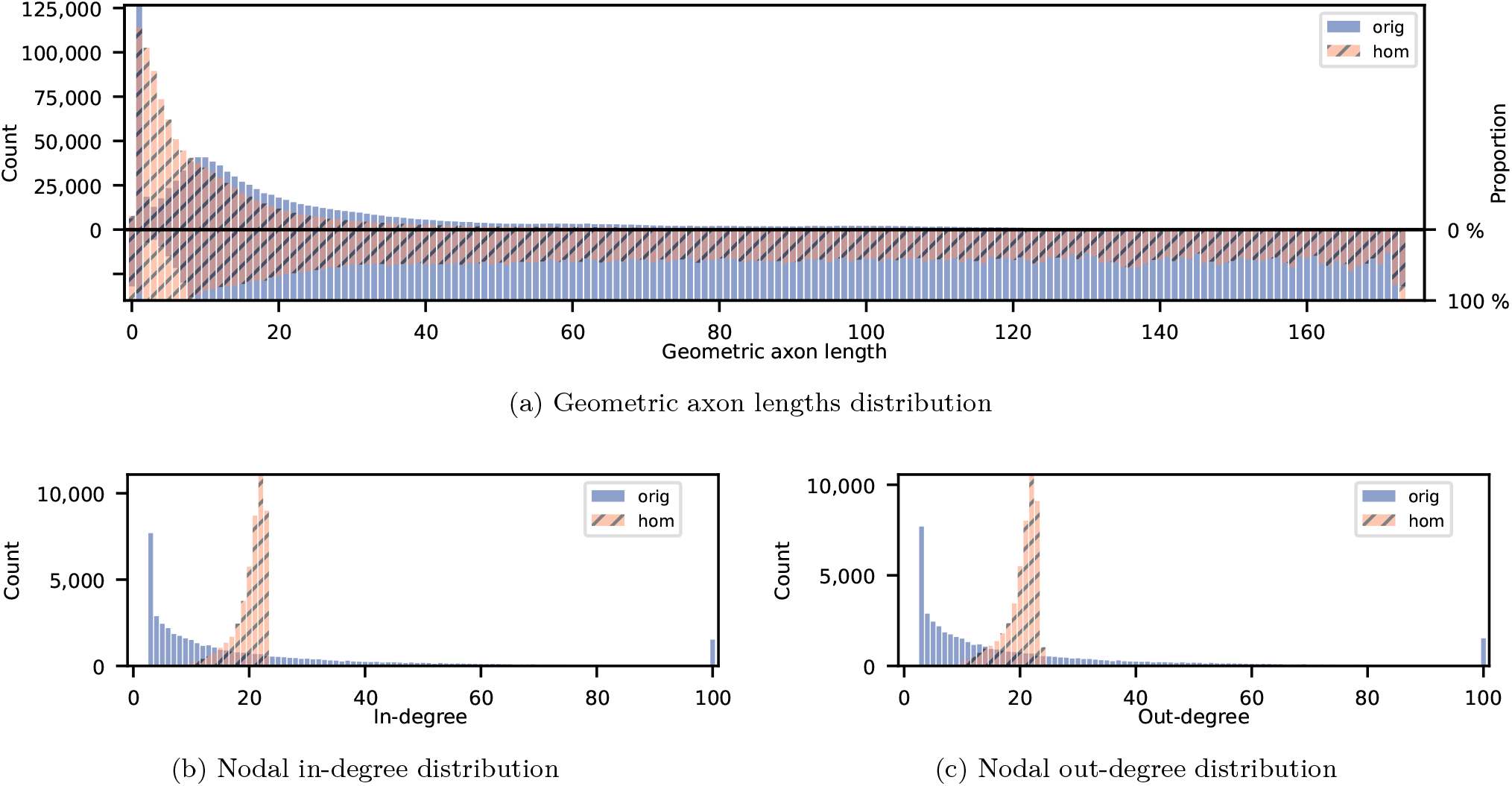
Effects of homeostatization on geometric-axon-length and node-degree distributions of a single connectome example. The distributions of the original connectome (i.e., freshly neuronized) are labeled *orig*, while the distributions of the homeostatized connectome are labeled *hom*. The distributions appear in solid blue and hatched orange, respectively; the hatched orange distribution is plotted on top of the solid blue distribution. Geometric axon length is the Euclidean distance between pre- and postsynaptic neurons. The upper panel of the histogram (left y-axis) shows the absolute count of each length bin, while the lower panel (right y-axis) shows the relative frequency (percent). For each bin, the maximum axon-length count from either the original or the homeostatized connectome, whichever is larger, defines 100 %. The percentage proportion of the maximum count is shown for the other. Comparing the percentage values of axon-length counts before and after homeostatization reveals that homeostatization preserves approximately 40 % of the original connectomes’ long-range connections while increasing the fraction of short-range connections. Homeostatization shifts the degrees from a fat-tailed distribution in the original connectome to a more normalized distribution with a cutoff to the right. This shift is a direct consequence of homeostatic synapse formation and deletion and is unavoidable with a common activity set-point and binary weights. Neurons simply cannot afford higher in-degrees because this would carry the activation above the set-point and lead to immediate synapse breakdown. Lower in-degrees are possible because some neurons have not yet reached the activity set-point. The out-degree distribution follows the same trend, but it is slightly broader because the out-degree is only indirectly coupled with neuronal activity.

**Fig. A6:**
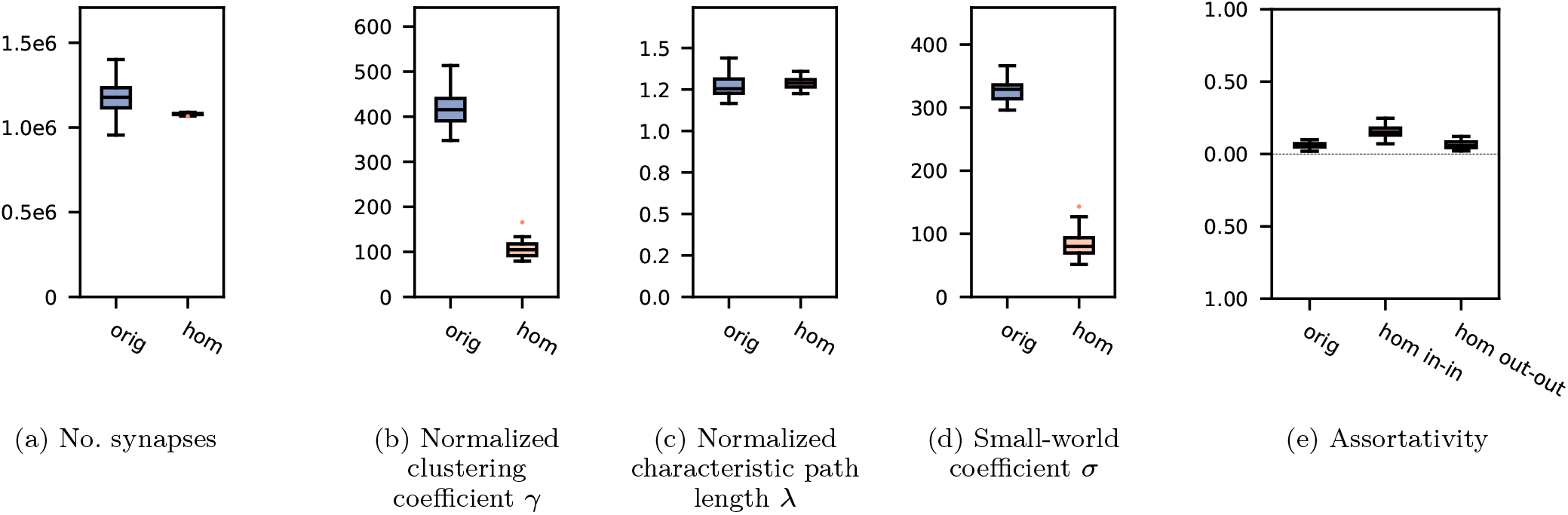
Effects of homeostatization on complex network measures [67] across the entire population of 36 connectomes. The box plots illustrate the distribution of these measures across all 36 connectomes. The values of the original connectomes (i.e., freshly neuronized), labeled *orig*, are shown in blue, and the values of the homeostatized connectomes, labeled *hom*, are shown in orange. The box plots include the median (black line), the interquartile range (IQR, box), the whiskers (±1.5 IQR), and the outliers (colored dots). For assortativity, we present one distribution for the unidirectional, original connectomes and two for the bidirectional, homeostatized ones, one for incoming connections and one for outgoing connections. This is because DTI yields only undirected connections, which homeostatization converts to directed ones. The slightly increased correlation values for the in-degree assortativity reflect the narrower degree distributions after homeostatization.

## Footnotes

1 https://fsl.fmrib.ox.ac.uk/fsl/fslwiki

2 https://fsl.fmrib.ox.ac.uk/fsl/docs/diffusion/topup

3 http://dsi-studio.labsolver.org

4 https://github.com/tuda-hpclab/relearn

5 https://github.com/tuda-hpclab/neurograph

## Notes

### Competing Interest Statement

The authors have declared no competing interest.

### Summary of Updates

This revision presents the computational workflow more clearly, corrects several inaccuracies in the original posting, and improves the overall clarity of the manuscript.

## References

[1] S. Herculano-Houzel, “The remarkable, yet not extraordinary, human brain as a scaled-up primate brain and its associated cost,” in In the Light of Evolution VI: Brain and Behavior, G. F. Striedter, J. C. Avise, and F. J. Ayala, Eds. Washington, DC: National Academies Press, 2013, ch. 8.

[2] M. Kaiser, Changing Connectomes: Evolution, Development, and Dynamics in Network Neuroscience. Cambridge, MA: MIT Press, 2020.

[3] G. F. Striedter, Principles of Brain Evolution. Sunderland, MA: Sinauer Associates, 2005.

[4] F. A. Azevedo, L. R. Carvalho, L. T. Grinberg, J. M. Farfel, R. E. Ferretti, R. E. Leite, W. J. Filho, R. Lent, and S. Herculano-Houzel, “Equal numbers of neuronal and nonneuronal cells make the human brain an isometrically scaled-up primate brain,” Journal of Comparative Neurology, vol. 513, no. 5, pp. 532–541, 2009.

[5] B. Pakkenberg, D. Pelvig, L. Marner, M. J. Bundgaard, H. J. G. Gundersen, J. R. Nyengaard, and L. Regeur, “Aging and the human neocortex,” Experimental Gerontology, vol. 38, no. 1, pp. 95–99, 2003.

[6] M. Cao, H. Huang, and Y. He, “Developmental connectomics from infancy through early childhood,” Trends in Neurosciences, vol. 40, no. 8, pp. 494–506, 2017.

[7] S. H. Bennett, A. J. Kirby, and G. T. Finnerty, “Rewiring the connectome: Evidence and effects,” Neuroscience & Biobehavioral Reviews, vol. 88, pp. 51–62, 2018.

[8] C. Lee, B. H. Lee, H. Jung, C. Lee, Y. Sung, H. Kim, J. Kim, J. Y. Shim, J.-i. Kim, D. I. Choi et al., “Hippocampal engram networks for fear memory recruit new synapses and modify preexisting synapses in vivo,” Current Biology, vol. 33, no. 3, pp. 507–516, 2023.

[9] D. B. Chklovskii, B. W. Mel, and K. Svoboda, “Cortical rewiring and information storage,” Nature, vol. 431, no. 7010, pp. 782–788, 2004.

[10] I. E. Dammasch, G. P. Wagner, and J. R. Wolff, “Self-stabilization of neuronal networks. I. The compensation algorithm for synaptogenesis,” Biological Cybernetics, vol. 54, no. 4–5, pp. 211–222, 1986.

[11] I. E. Dammasch, G. P. Wagner, and J. R. Wolff, “Self-stabilization of neuronal networks. II. Stability conditions for synaptogenesis,” Biological Cybernetics, vol. 58, no. 3, pp. 149–158, 1988.

[12] M. Butz and A. van Ooyen, “A simple rule for dendritic spine and axonal bouton formation can account for cortical reorganization after focal retinal lesions,” PLoS Computational Biology, vol. 9, no. 10, p. e1003259, 2013.

[13] I. E. Dammasch, “Structural realization of a Hebb-type learning rule,” in Models of Brain Function, R. M. J. Cotterill, Ed. Cambridge: Cambridge University Press, 1989, pp. 539–552.

[14] J. V. Gallinaro, N. Gašparović, and S. Rotter, “Homeostatic control of synaptic rewiring in recurrent networks induces the formation of stable memory engrams,” PLOS Computational Biology, vol. 18, no. 2, p. e1009836, 2022.

[15] M. Kaster, F. Czappa, M. Butz-Ostendorf, and F. Wolf, “Building a realistic, scalable memory model with independent engrams using a homeostatic mechanism,” Frontiers in Neuroinformatics, vol. 18, p. 1323203, 2024.

[16] R. Quian Quiroga, “Concept cells: the building blocks of declarative memory functions,” Nature Reviews Neuroscience, vol. 13, pp. 587–597, 2012.

[17] R. Quian Quiroga, “Searching for the neural correlates of human intelligence,” Current Biology, vol. 30, no. 8, pp. R335–R338, 2020.

[18] L. Reddy and S. J. Thorpe, “Concept cells through associative learning of high-level representations,” Neuron, vol. 84, no. 2, pp. 248–251, 2014.

[19] R. Quian Quiroga, A. Kraskov, F. Mormann, I. Fried, and C. Koch, “Single-cell responses to face adaptation in the human medial temporal lobe,” Neuron, vol. 84, no. 2, pp. 363–369, 2014.

[20] R. Quian Quiroga, A. Kraskov, C. Koch, and I. Fried, “Explicit encoding of multimodal percepts by single neurons in the human brain,” Current Biology, vol. 19, no. 15, pp. 1308–1313, 2009.

[21] M. Ison, R. Quian Quiroga, and I. Fried, “Fast remapping of single neuron responses in the human medial temporal lobe,” in Society for Neuroscience Abstracts, vol. 279, no. 15, Nov. 2010, p. 72.

[22] R. Paz, H. Gelbard-Sagiv, R. Mukamel, M. Harel, R. Malach, and I. Fried, “A neural substrate in the human hippocampus for linking successive events,” Proceedings of the National Academy of Sciences, vol. 107, no. 13, pp. 6046–6051, 2010.

[23] D. Marr, “Simple memory: a theory for archicortex,” Philosophical Transactions of the Royal Society of London. Series B: Biological Sciences, vol. 262, no. 841, pp. 23–81, 1971.

[24] D. O. Hebb, The Organization of Behavior: A Neuropsychological Theory. New York: John Wiley & Sons, 1949.

[25] M. F. Bear and R. C. Malenka, “Synaptic plasticity: LTP and LTD,” Current Opinion in Neurobiology, vol. 4, no. 3, pp. 389–399, 1994.

[26] R. C. Malenka and M. F. Bear, “LTP and LTD: an embarrassment of riches,” Neuron, vol. 44, no. 1, pp. 5–21, 2004.

[27] E. T. Rolls, “Roles of LTP and LTD in neuronal network in the brain,” in Cortical Plasticity: From Synapses to Maps, D. E. Feldman, Ed. New York: Garland Science, 2020, pp. 223–250.

[28] N. Caporale and Y. Dan, “Spike timing-dependent plasticity: a Hebbian learning rule,” Annual Review of Neuroscience, vol. 31, pp. 25–46, 2008.

[29] M. Butz, I. D. Steenbuck, and A. van Ooyen, “Homeostatic structural plasticity can account for topology changes following deafferentation and focal stroke,” Frontiers in Neuroanatomy, vol. 8, p. 115, 2014.

[30] X. Chen, Y. Wang, S. J. Kopetzky, M. Butz-Ostendorf, and M. Kaiser, “Connectivity within regions characterizes epilepsy duration and treatment outcome,” Human Brain Mapping, vol. 42, no. 12, pp. 3777–3791, 2021.

[31] H. Axer, S. Beck, M. Axer, F. Schuchardt, J. Heepe, A. Flücken, M. Axer, A. Prescher, and O. W. Witte, “Microstructural analysis of human white matter architecture using polarized light imaging: Views from neuroanatomy,” Frontiers in Neuroinformatics, vol. 5, p. 28, 2011.

[32] R. M. Willems and K. Henke, “Imaging human engrams using 7 Tesla magnetic resonance imaging,” Hippocampus, vol. 31, no. 12, pp. 1257–1270, 2021.

[33] P. Sanz-Leon, S. A. Knock, M. M. Woodman, L. Domide, J. Mersmann, A. R. McIntosh, and V. K. Jirsa, “The Virtual Brain: a simulator of primate brain network dynamics,” Frontiers in Neuroinformatics, vol. 7, p. 10, 2013.

[34] R. S. Desikan, F. Ségonne, B. Fischl, B. T. Quinn, B. C. Dickerson, D. Blacker, R. L. Buckner, A. M. Dale, R. P. Maguire, B. T. Hyman et al., “An automated labeling system for subdividing the human cerebral cortex on MRI scans into gyral based regions of interest,” NeuroImage, vol. 31, no. 3, pp. 968–980, 2006.

[35] J. O’Keefe and J. Dostrovsky, “The hippocampus as a spatial map: preliminary evidence from unit activity in the freely-moving rat,” Brain Research, vol. 34, no. 1, pp. 171–175, 1971.

[36] M. K. Benna and S. Fusi, “Place cells may simply be memory cells: memory compression leads to spatial tuning and history dependence,” Proceedings of the National Academy of Sciences, vol. 118, no. 51, p. e2018422118, 2021.

[37] P. Baraduc, J.-R. Duhamel, and S. Wirth, “Schema cells in the macaque hippocampus,” Science, vol. 363, no. 6427, pp. 635–639, 2019.

[38] R. Quian Quiroga, L. Reddy, G. Kreiman, C. Koch, and I. Fried, “Invariant visual representation by single neurons in the human brain,” Nature, vol. 435, no. 7045, pp. 1102–1107, 2005.

[39] S. A. Josselyn and S. Tonegawa, “Memory engrams: recalling the past and imagining the future,” Science, vol. 367, no. 6473, p. eaaw4325, 2020.

[40] R. Semon, Die Mneme: Als erhaltendes Prinzip im Wechsel des organischen Geschehens. Leipzig: Wilhelm Engelmann, 1904.

[41] M. Geva-Sagiv, E. A. Mankin, D. Eliashiv, S. Epstein, N. Cherry, G. Kalender, N. Tchemodanov, Y. Nir, and I. Fried, “Augmenting hippocampal–prefrontal neuronal synchrony during sleep enhances memory consolidation in humans,” Nature Neuroscience, vol. 26, no. 6, pp. 1100–1110, 2023.

[42] T. Xu, X. Yu, A. J. Perlik, W. F. Tobin, J. A. Zweig, K. Tennant, T. Jones, and Y. Zuo, “Rapid formation and selective stabilization of synapses for enduring motor memories,” Nature, vol. 462, pp. 915–919, 2009.

[43] T. Kitamura, S. K. Ogawa, D. S. Roy, T. Okuyama, M. D. Morrissey, L. M. Smith, R. L. Redondo, and S. Tonegawa, “Engrams and circuits crucial for systems consolidation of a memory,” Science, vol. 356, no. 6333, pp. 73–78, 2017.

[44] P. Lavenex and D. G. Amaral, “Hippocampal–neocortical interaction: a hierarchy of associativity,” Hippocampus, vol. 10, no. 4, pp. 420–430, 2000.

[45] J.-H. Choi, S.-E. Sim, J.-i. Kim, D. I. Choi, J. Oh, S. Ye, J. Lee, T. Kim, H.-G. Ko, C.-S. Lim, and B.-K. Kaang, “Interregional synaptic maps among engram cells underlie memory formation,” Science, vol. 360, no. 6387, pp. 430–435, 2018.

[46] D. S. Roy, Y.-G. Park, M. E. Kim, Y. Zhang, S. K. Ogawa, N. DiNapoli, X. Gu, J. H. Cho, H. Choi, L. Kamentsky, J. Martin, O. Mosto, T. Aida, K. Chung, and S. Tonegawa, “Brain-wide mapping reveals that engrams for a single memory are distributed across multiple brain regions,” Nature Communications, vol. 13, no. 1, p. 1799, 2022.

[47] C. Du, K. Fu, B. Wen, Y. Sun, J. Peng, W. Wei, Y. Gao, S. Wang, C. Zhang, J. Li, S. Qiu, L. Chang, and H. He, “Human-like object concept representations emerge naturally in multimodal large language models,” Nature Machine Intelligence, vol. 7, no. 6, pp. 860–875, 2025.

[48] C. Gastaldi, T. Schwalger, E. De Falco, R. Quian Quiroga, and W. Gerstner, “When shared concept cells support associations: theory of overlapping memory engrams,” PLOS Computational Biology, vol. 17, no. 12, p. e1009691, 2021.

[49] R. Hecht-Nielsen, Confabulation Theory: The Mechanism of Thought. Springer, 2007.

[50] D. J. Amit, H. Gutfreund, and H. Sompolinsky, “Storing infinite numbers of patterns in a spin-glass model of neural networks,” Physical Review Letters, vol. 55, no. 14, pp. 1530–1533, 1985.

[51] A. Knoblauch, “Impact of structural plasticity on memory formation and decline,” in The Rewiring Brain, A. van Ooyen and M. Butz-Ostendorf, Eds. San Diego: Academic Press, 2017, pp. 361–386.

[52] J. N. Bourne and K. M. Harris, “Coordination of size and number of excitatory and inhibitory synapses results in a balanced structural plasticity along mature hippocampal CA1 dendrites during LTP,” Hippocampus, vol. 21, no. 4, pp. 354–373, Apr. 2011.

[53] J.-P. Changeux and A. Danchin, “Selective stabilisation of developing synapses as a mechanism for the specification of neuronal networks,” Nature, vol. 264, no. 5588, pp. 705–712, 1976.

[54] A. Whittall, “Leitmotif,” Grove Music Online, 2001.

[55] D. R. Hofstadter, “Church, Turing, Tarski, and others,” in Gödel, Escher, Bach: An Eternal Golden Braid. New York: Basic Books, 1979.

[56] E. M. Izhikevich and G. M. Edelman, “Large-scale model of mammalian thalamocortical systems,” Proceedings of the National Academy of Sciences, vol. 105, no. 9, pp. 3593–3598, 2008.

[57] C. Eliasmith, T. C. Stewart, X. Choo, T. Bekolay, T. DeWolf, Y. Tang, and D. Rasmussen, “A large-scale model of the functioning brain,” Science, vol. 338, no. 6111, pp. 1202–1205, 2012.

[58] H. E. Wang, P. Triebkorn, M. Breyton, B. Dollomaja, J.-D. Lemarechal, S. Petkoski, P. Sorrentino, D. Depannemaecker, M. Hashemi, and V. K. Jirsa, “Virtual brain twins: from basic neuroscience to clinical use,” National Science Review, vol. 11, no. 5, p. nwae079, 2024.

[59] F. Hutchings, C. E. Han, S. S. Keller, B. Weber, P. N. Taylor, and M. Kaiser, “Predicting surgery targets in temporal lobe epilepsy through structural connectome based simulations,” PLOS Computational Biology, vol. 11, no. 12, p. e1004642, 2015.

[60] Q. Fang and D. A. Boas, “Tetrahedral mesh generation from volumetric binary and grayscale images,” in 2009 IEEE International Symposium on Biomedical Imaging: From Nano to Macro, 2009, pp. 1142–1145.

[61] P. N. Taylor, Y. Wang, and M. Kaiser, “Within brain area tractography suggests local modularity using high resolution connectomics,” Scientific Reports, vol. 7, p. 39859, 2017.

[62] E. M. Izhikevich, “Simple model of spiking neurons,” IEEE Transactions on Neural Networks, vol. 14, no. 6, pp. 1569–1572, 2003.

[63] B. A. Richards and K. P. Kording, “The study of plasticity has always been about gradients,” The Journal of Physiology, vol. 601, no. 15, pp. 3141–3149, 2023.

[64] V. De Paola, A. Holtmaat, G. Knott, S. Song, L. Wilbrecht, P. Caroni, and K. Svoboda, “Cell type-specific structural plasticity of axonal branches and boutons in the adult neocortex,” Neuron, vol. 49, no. 6, pp. 861–875, 2006.

[65] S. Rinke, M. Butz-Ostendorf, M.-A. Hermanns, M. Naveau, and F. Wolf, “A scalable algorithm for simulating the structural plasticity of the brain,” Journal of Parallel and Distributed Computing, vol. 120, pp. 251–266, 2018.

[66] F. Czappa, A. Geiß, and F. Wolf, “Simulating structural plasticity of the brain more scalable than expected,” Journal of Parallel and Distributed Computing, vol. 171, pp. 24–27, 2023.

[67] M. Rubinov and O. Sporns, “Complex network measures of brain connectivity: uses and interpretations,” NeuroImage, vol. 52, no. 3, pp. 1059–1069, 2010.

